# Direct Thalamic Inputs to Hippocampal CA1 Transmit a Signal That Suppresses Ongoing Contextual Fear Memory Retrieval

**DOI:** 10.1101/2023.03.27.534420

**Authors:** Heather C. Ratigan, Seetha Krishnan, Shai Smith, Mark E. J. Sheffield

## Abstract

Memory retrieval of fearful experiences is essential for survival but can be maladaptive if not appropriately suppressed. Fear memories can be acquired through contextual fear conditioning (CFC) which relies on the hippocampus. The thalamic subregion Nucleus Reuniens (NR) is necessary for contextual fear extinction and strongly projects to hippocampal subregion CA1. However, the NR-CA1 pathway has not been investigated during behavior, leaving unknown its role in contextual fear memory retrieval. We implement a novel head-restrained virtual reality CFC paradigm and show that inactivation of the NR-CA1 pathway prolongs fearful freezing epochs, induces fear generalization, and delays extinction. We use *in vivo* sub-cellular imaging to specifically record NR-axons innervating CA1 before and after CFC. We find NR-axons become selectively tuned to freezing only after CFC, and this activity is well-predicted by an encoding model. We conclude that the NR-CA1 pathway actively suppresses fear responses by disrupting ongoing hippocampal-dependent contextual fear memory retrieval.

## INTRODUCTION

Flexibly encoding and retrieving memories of fearful events is a critically conserved survival behavior, as a single failure can be deadly. However, failing to suppress inappropriate fear responses can also have devastating consequences, manifesting as negative affective states in generalized anxiety disorder and post-traumatic stress disorder^1,2^. One way in which fear memories can be studied in the laboratory is through contextual fear conditioning (CFC), in which a spatial context, the conditioned stimulus (CS), is repeatedly paired with a noxious unconditioned stimulus (US), generally a mild shock^3–56^. Freezing is a species-specific fear response, and a quantifiable readout of contextual fear memory retrieval (CFMR) of the learned association^7,8^. With continued exposure to the CS in the absence of the US, freezing generally decreases and exploratory behavior increases - a process termed fear extinction. Fear extinction occurs as animals learn over time that the context no longer predicts shocks^9–12^. For extinction to occur, mice must therefore suppress CFMR during each fearful freezing epoch to avoid excessive freezing, which would be detrimental to survival. Therefore, mechanisms must exist in the brain to suppress CFMR as it is occurring.

Contextual fear in both mice and humans relies on coordinated brain regions including the Medial Prefrontal Cortex (mPFC), Thalamus, Amygdala, and Hippocampus^13^. The contextual component of these memories relies on the hippocampus, which retrieves and updates contextual fear memories^6,14–19^. Experimental inhibition of a subset of hippocampal neurons tagged using immediate early genes active during CFC is sufficient to suppress CFMR^20–22^. This suggests that natural suppression of ongoing CFMR must involve a circuit that can modulate hippocampal activity.

One potential source of this modulation is the ventral midline thalamic subregion, Nucleus Reuniens (NR). Sometimes termed ‘limbic thalamus’ for its diverse set of inputs from limbic-related regions in the brainstem, hypothalamus, amygdala, basal forebrain, mPFC, entorhinal cortex (EC), and hippocampal subregion CA1, NR sits at the nexus of emotional regulation and serves as a major communication hub among these limbic-activated areas^23–28^. While mPFC does not have a direct excitatory projection to CA1, it does send a strong excitatory projection to NR^23,24^.

The mPFC-NR projection and NR itself is necessary for both fear extinction and for preventing fear generalization to a neutral context, a process in which mice fail to form context-specific memory and additionally associate a non-shocked context with fear^29–35^. NR stimulation reduces contextual fear-induced immediate early gene expression in both mPFC and CA1^35,36^. While the roles of the mPFC-NR pathway and NR itself have been explored during CFMR, the role of the NR-CA1 pathway is unknown. We hypothesize that NR transmits a signal to CA1 to suppress ongoing CFMR, thereby reducing fear responses (freezing) and promoting exploratory behavior (movement).

To test our hypothesis, we used a chemogenetic approach to directly inhibit the NR-CA1 pathway, and 2-photon calcium imaging in head-restrained mice to record NR-axons in CA1, before and after CFC. While CFC induction in head-restrained mice in virtual reality (VR) has been attempted, none to our knowledge have replicated the characteristic ‘freezing’ behavior of freely-moving mice in real-world CFC^37,38^. We therefore developed a new VR-based CFC paradigm (VR-CFC), using a conductive fabric to deliver mild tail shocks that induces context-dependent freezing behavior in mice. By combining VR-CFC, targeted chemogenetic NR-CA1 inactivation, and 2-photon NR-axonal calcium imaging, we were able to determine the role of the NR-CA1 pathway in CFMR.

## RESULTS

### Contextual Fear Conditioning and Extinction in Virtual Contexts

To ensure mice were comfortable with the VR setup before shocks were delivered, we trained water-restricted mice to run in a VR context for water rewards until they reached ~4 traversals of the context per minute, as previously discussed^39,40^. To avoid confounds from the water reward during CFC, these trained mice were then introduced to two novel VR contexts without a water reward. At this stage, the custom-designed conductive tailcoat was fitted to their tails (Fig. 1A, Extended Data Fig. 1A; Method Details). Mice spent ~5 minutes in each novel VR context which allowed them to habituate to running with the tailcoat. (Fig. 1A, Extended Data Fig. 1A; Method Details).

**Figure 1.**
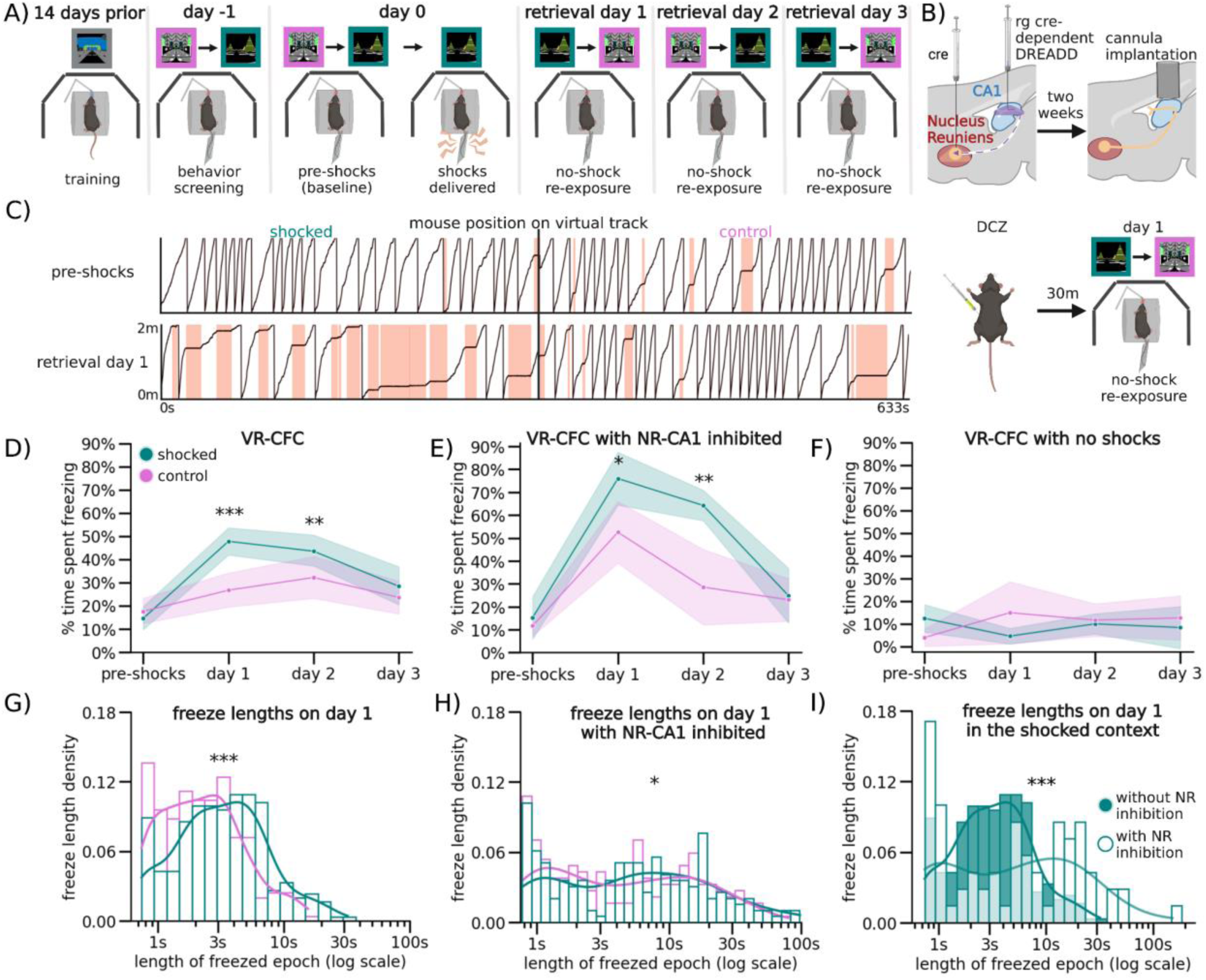
Nucleus Reuniens-CA1 Pathway Inhibition During Memory Retrieval Increases Fearful Behavior. (A) After training, head-restrained mice were put in two unrewarded contexts. They then received mild tail shocks in only one context (shocked), and not the other (control). They were re-exposed to both contexts in pseudo-random order for 3 retreival days. (B) Top: NR-CA1 axons were inhibited using cre-dependent DREADDs, resulting in a subset of CA1-projecting NR neurons expressing HM4di receptors. Bottom: ~30 minutes before day 1 context re-exposure, mice received 0.1 mg/kg of the HM4d agonist DCZ. (C) Example track position from a mouse on day 0 pre-shocks (Top), and on retrieval day 1 (Bottom), in the shocked (left) and control (right) contexts. Red shading indicates freezing epochs. (D) Line indicates mean, shading indicates 95% CI. Mice froze significantly more post-shocks, and froze more in the shocked context (teal) than the control context (pink), on both retrieval days 1 and 2. By retrieval day 3, freezing between contexts was equivalent (N = 20 mice; Wilcoxon Rank Sum, P=pre-shocks: 1.00, day 1: 4.12e-4, day 2: 9.87e-3, day 3: 0.55). (E) NR-CA1 inhibited mice froze significantly more in both contexts than NR-CA1 intact mice (D) on retrieval day 1 (N = 5 mice). Freezing in these mice remained elevated in the shocked, but not the control, context on day 2. Freezing levels in both contexts became similar by day 3. (Wilcoxon Rank Sum, P = pre-shocks: 0.76, day 1: 1.43e-30, day 2: 8.11e-3, day 3: 1.00). (F) Mice were otherwise trained as in Fig. 1A, but not shocked on day 0. These mice froze at baseline levels across all days, (N = 4 mice, Wilcoxon Rank Sum, P = pre-shocks: 0.13, day 1: 0.5, day 2: 0.5, day 3: 0.25). (G) Kernel density estimates and density histogram of freeze epochs were shorter in the control than the shocked context on day 1 (Mann-Whitney U, P = 4.30e-5). (H) Freeze epochs were similar during NR inhibition relative to day 1 in NR-CA1 intact mice (Mann-Whitney U, P = Shocked: 3.63e-4, Control: 4.22e-4). (I) Freeze epochs in the shocked context with NR inhibition skewed longer compared to without inhibition on day 1 (Mann-Whitney U, P = 2.16e-6).

Mice in all experimental conditions that continued to meet the criterion for movement (i.e. >4 traversals per minute) were advanced to the next stage the following day (36/64 mice), where they were re-exposed to both novel contexts for ~5 minutes each, then were administered 6 mild 0.6 mA tail shocks through the tailcoat for a duration of 1 s each, 20-26 s apart, (Fig. 1A: day 0). These shocks were delivered at pseudorandom locations throughout one of the contexts (‘shocked’), but not the other (‘control’; Extended Data Fig. 1B). Mice responded to each tail shock with an abrupt stereotyped increase in running speed, a behavioral validation of successful shock delivery (Extended Data Fig. 1D). To test for CFMR and subsequent fear memory extinction, mice were then re-exposed for ~5 minutes each, in a pseudorandom order, to both the shocked and control contexts while wearing the tailcoat for the following three retrieval days (Fig. 1A: day 1-3).

On retrieval day 1, the subset of mice used for VR-CFC (N = 20) froze in the control context on average 26.7 ± 7.2% of the time (95% CI), while mice in the shocked context froze significantly more, on average 47.8 ± 8.3% of the time (Fig. 1D: day 1, Wilcoxon Rank Sum shocked versus control context, P = day 0 pre-shocks: 0.0076, day 1: 0.0014, day 2: 0.0009 day 3: 1.00). Compared to baseline levels of freezing pre-shocks, freezing on retrieval day 1 in both contexts was elevated, although mice increased their freezing significantly more in the shocked context than in the control context (P = Shocked: 5.04e-07, Control: 0.0021, Extended Data Fig. 2B). This remained true on retrieval day 2, as mice continued to freeze at significantly elevated levels of 42.7 ± 8.0% in the shocked context compared to 28.3 ± 7.1% in the control context, with freezing levels in both contexts significantly higher than pre-shock freezing levels (Fig. 1D: day 2; Extended Data Fig. 2B: day 2, P=Shocked: 3.89e-07, Control: 0.00191). By the third day of retrieval, mice froze at similar levels across contexts, at 28.4 ± 9.6% in the shocked versus 23.1 ± 7.2% in the control context, and returned to near baseline in the shocked context and baseline in the control context (Extended Data Fig. 2B: day 3).

**Figure 2.**
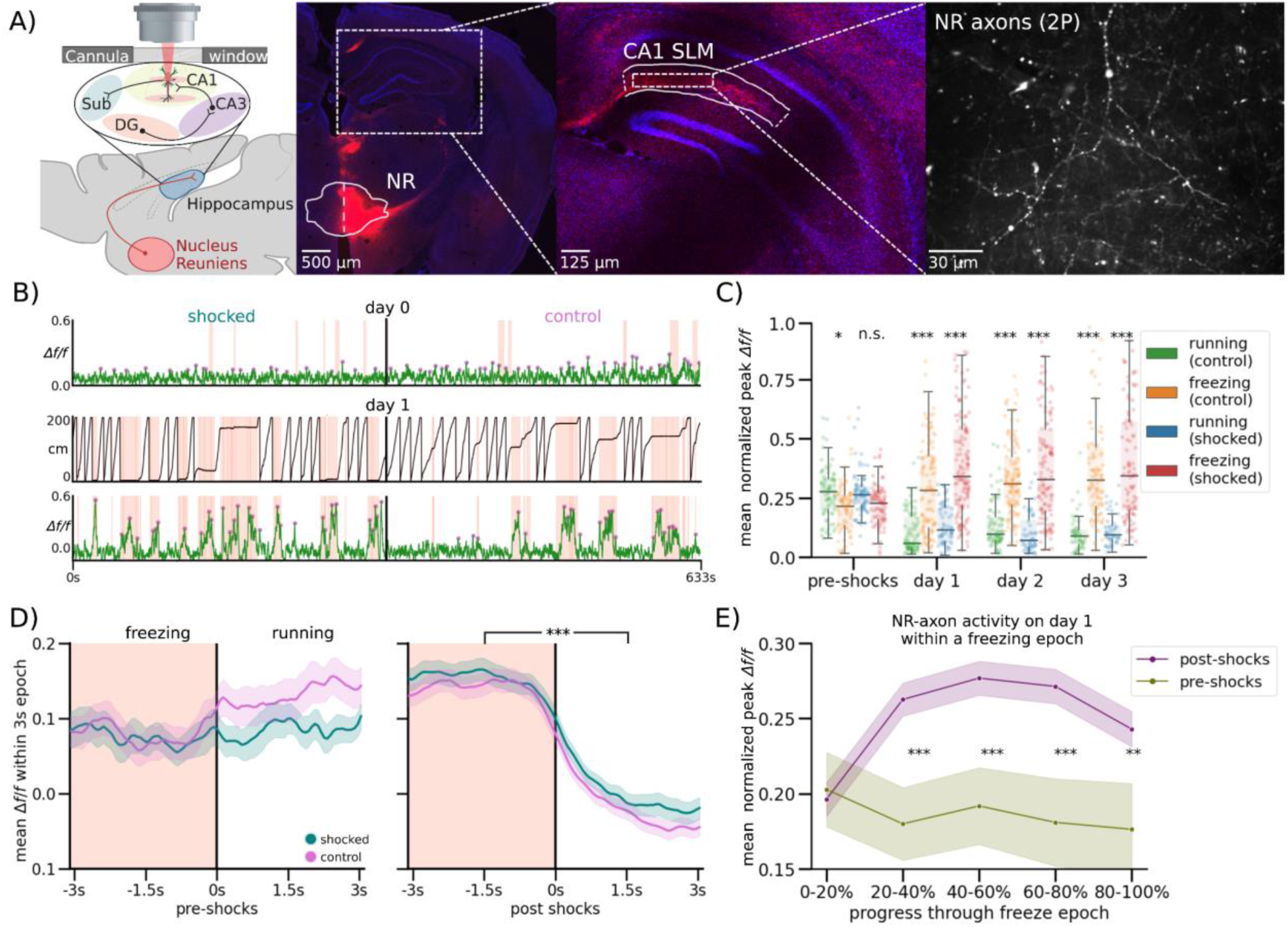
Nucleus Reuniens-CA1 Axons Increase Activity During Fearful Behavior Following CFC. (A) Left: Schematic representation of NR axonal imaging. Mice were trained as in Fig. 1A. Middle left: NR mRuby expression in Nucleus Reuniens under confocal imaging. Middle right: Axonal expression in the hippocampus limited to the SLM layer in subiculum and CA1. Right: Example average FOV of NR axons in SLM through 2-photon during mouse behavior. (B) Example mouse NR axonal activity pre-shocks (Top) and on day 1 (Bottom) in the shocked (left) and control (right) contexts. Red shading indicates freezing epochs. Middle trace is the mouse position on retrieval day 1. (C) Normalized mean *Δf/f* of axonal peaks per freezing epoch plotted as dots, boxplot indicates median, 25-75th interquartile range, whiskers include all data points not determined to be outliers. In both contexts, mean normalized axonal activity increases post-shocks compared to pre-shocks, and remains elevated during post-shocks freezing epochs as compared to running epochs. Correspondingly, activity in running epochs decreased post-shocks from pre-shocks (Student’s T, P = pre-shocks: 0.085, 0.039, day 1: 9.42e-15, 6.27e-13, day 2: 6.64e-18, 7.19e-13, day 3, 1.22e-12, 1.47e-13). (D) Peach shading (left) indicates freezing epochs. Line indicates mean, shading indicates 95% CI. Pre-shocks, NR-CA1 pathway axonal activity is comparable during freezing and running epochs. Post-shocks, activity is significantly elevated during freezing when compared to running epochs (Wilcoxon Rank Sum left: P = 0.14, right: P = 1.37e-52). Freezing epochs displayed are 3-4 s long, additional epoch windows are shown in Extended Data Fig 4F). (E) Normalized mean *Δf/f* of axonal peaks were binned into 5 categories based on percent progress throughout the freezing epoch. For the majority of the pause (20-100% freeze progress) in both contexts (purple), axonal activity was significantly increased than activity before shocks (green) (Student’s T, P = 0-20%: 0.81, 20-40%: 1.71e-3, 40-60%: 9.75e-4, 60-80%: 1.46e-3, 80-100%: 8.70e-3).

While freezing quantity differed between contexts and days of retrieval, freezing position was distributed evenly across all track locations in both contexts on all retrieval days. This shows that mice associated fear with the entire context, and not specific locations along the track or near specific objects in VR (Extended Data Fig. 1C). As an additional control, a separate group of mice went through the same process but were never shocked in either context. These mice froze significantly less on retrieval days 1-3 (on average 11.6 ± 8.7%) without any significant differences to freezing on day 0 or between contexts (Fig. 1F, Extended Data Fig. 2H). These spontaneous freezing events in both the control condition and the pre-shocked contexts (i.e. before the delivery of any shocks) could potentially be caused by the lack of water reinforcement, the presence of the tail coat itself, or an unrelated temporary disinterest in running, and provide a within-mouse comparison to post-shock fear-evoked freezing.

To further quantify freezing behavior, we measured the duration of each individual freezing event (freezing epoch) and found that freezing epochs were longer in the shocked versus the control context on retrieval day 1 (Fig. 1G). Freezing epochs remained longer on day 2 in the shocked compared to the control context, however, they became similar by day 3 (Extended Data Fig. 2F-I), corresponding with the total time spent freezing. Our results show that VR-CFC produces robust CFMR, that can be measured via context-specific increase in freezing and can be reliably extinguished following ~3 days of re-exposure to the shocked context in the absence of additional shocks.

### Inhibition of the NR-CA1 Pathway during CFMR Increases Freezing Behavior

To test the involvement of NR-CA1 projecting neurons in CFMR, we designed a designer receptor exclusively activated by designer drugs (DREADD) based inhibition paradigm^41–43^ (Fig. 1b). We injected a Cre-expressing virus bilaterally in NR, and a retrograde Cre-dependent virus carrying the inhibitory G(i)-coupled DREADD receptor, hM4Di-DREADD, bilaterally in the SLM of dorsal CA1 where hippocampal-projecting NR-axons terminate (Fig. 2A)^44^. This enabled us to intraperitoneally (IP) inject the hM4Di agonist, deschloroclozapine dihydrochloride (DCZ)^43^, before the first post-shock re-exposure to the contexts on retrieval day 1, therefore selectively inhibiting a subset of NR-CA1 projecting neurons during ongoing CFMR. To ensure our injection paradigm and administration of DCZ did not alter context-dependent fear behavior, we had two DREADD-control groups. One group (N = 4) expressed mCherry in place of hM4Di, and received DCZ on retrieval day 1. A separate group (N = 4) expressed the hM4Di receptor, and received saline instead of DCZ on retrieval day 1 (Extended Data Fig. 3A). In both control groups (Extended Data Fig. 3A), freezing behavior was similar to the experimental mice shown in Fig. 1D, and the groups were thus combined and termed the NR-CA1 intact group for further analysis. We then compared the behavioral impact of inhibiting the NR-CA1 pathway on day 1, and on subsequent retrieval days 2 and 3 with the NR-CA1 intact group (Fig. 1B Bottom).

**Figure 3.**
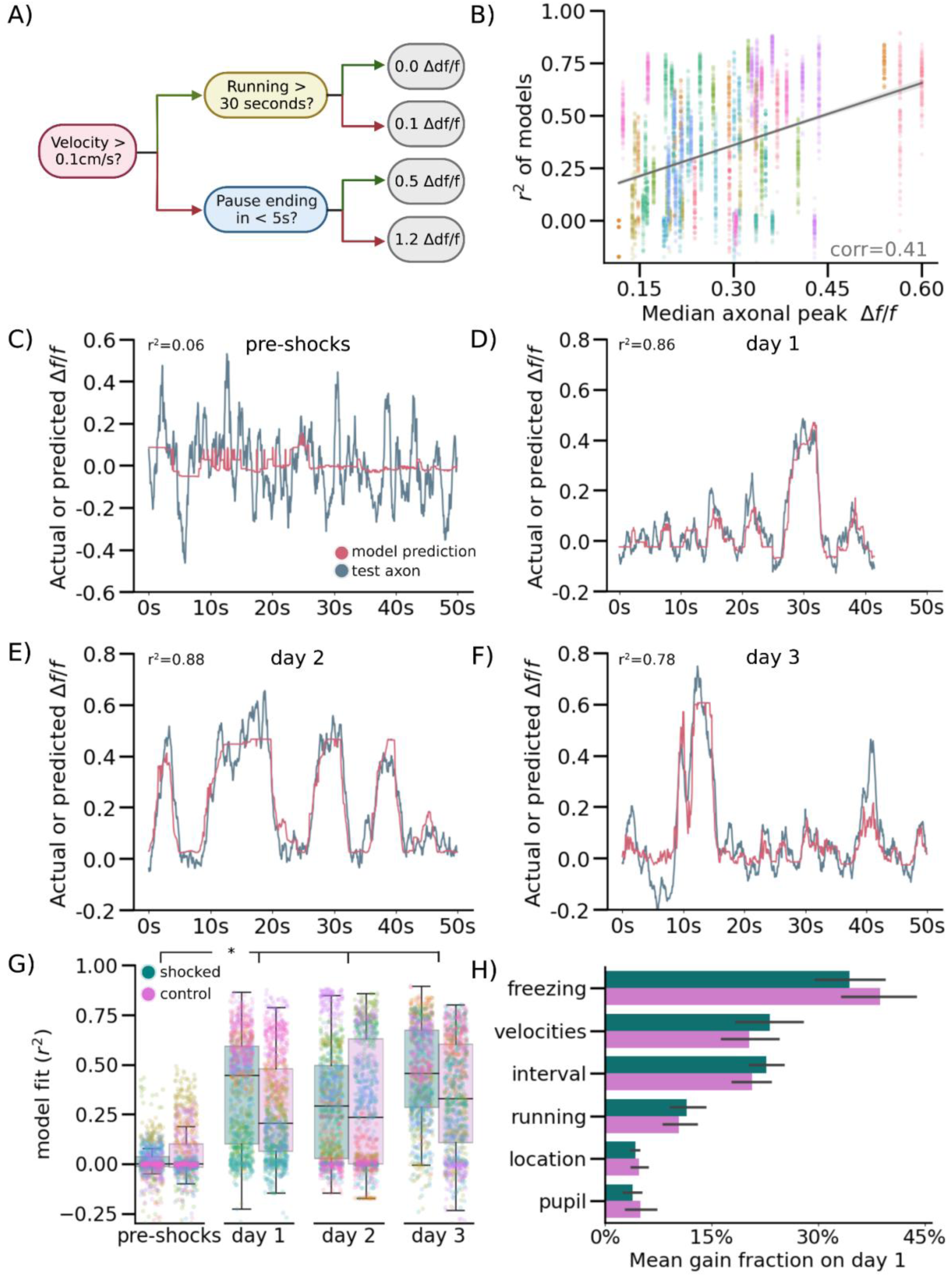
Nucleus Reuniens-CA1 Axon Activity is Accurately Predicted by a Computational Model Following CFC. (A) Simplified schematic of decision tree prediction. (B) Model performed better on axons with higher fluorescence signals. Goodness of model fit r^2^ was calculated for all model runs and plotted against median unnormalized axonal peak height as a proxy for data quality, color coded per mouse (N=8000 total runs; 10 mice, run 100 times per mouse, day, and context). (C-F) Examples of model prediction for the same axon in the same mouse tracked across days in the shocked context. Matched control context model examples are shown in Extended Data Fig. 6D). (G) Points indicate goodness of fit r^2^ for each model run, color coded by mouse, boxplot indicates median r^2^, 25-75th interquartile range, whiskers include all data points not determined to be outliers. Median model performance improved for both the shocked (pink) and control (teal) contexts across all days post-shocks, compared to pre-shock, in 8/10 imaged mice (Wilcoxon Rank Sum, P = Shocked: day 1: 3.10e-2, day 2: 2.46e-2, day 3: 1.85e-2, Control: day 1: 4.23e-2, day 2: 3.48e-2, day 3: 3.11e-2). (H) Mean gain fraction plotted per category, error bar indicates SEM. Model parameters pertaining to information about pausing, velocity, and duration of time paused or remaining in either a pausing or running interval (‘interval’) were used more than model parameters pertaining to running information, location on the track, or pupil information. Full gain fractions are shown in Extended Data Fig 6B.

We found that in the shocked context on retrieval day 1, NR-CA1 inhibited mice (N = 5) spent ~57% more time freezing than NR-CA1 intact mice (77.8% ± 12.4% versus 49.3% ± 10.1%; Fig. 1D and E, day 1; P=7.43e-07; see also Extended Data Fig. 3C). Time spent freezing during NR-CA1 inhibition on day 1 remained significantly higher in the shocked versus the control context (Fig. 1E; P = 0.0026), even though freezing levels also increased in the control context compared to NR-CA1 intact mice (Extended Data Fig. 3D). In both contexts, the dynamics of freezing behavior changed during NR-CA1 inhibition. Freezing epochs lengthened, as mice froze 284% longer on average in the NR-CA1 inhibited mice (21.1 s) compared to intact mice (5.5 s) in the shocked context (Fig. 1G versus 1H). There was a similar average increase in freezing lengths of 188% in the control context (12.1 s in inhibited vs 4.2 s in intact mice), albeit not as high as in the shocked context (Fig. 1G control context; Fig. 1H control context). These findings suggest that inhibiting the NR-CA1 pathway during CFMR increases the time spent in a contiguous, ongoing state of fear in both contexts as revealed by the lengthening of individual freezing epochs. This suggests that the NR-CA1 pathway may be suppressing CFMR in both appropriate (shocked) and inappropriate (unshocked control) contexts, to reduce fearful freezing.

Given these findings, we asked if mice could still discriminate well between the shocked and the control context after NR-CA1 inhibition. To do so, we calculated a discrimination index^29^ which revealed a significant decrease in discrimination between the shocked and control contexts on day 1 from 32.2% ± 6.6% with NR-CA1 intact to 18.1% ± 6.3% with NR-CA1 inhibited (Extended Data Fig. 4B, P = 0.0034), suggesting that inhibition of the NR-CA1 pathway reduces fear-induced contextual discrimination.

The long-term impact of the increase in fearful behavior on retrieval day 1 caused by NR-CA1 inhibition could be seen on retrieval day 2 where freezing levels remain higher than in NR-CA1 intact mice, even though the NR-CA1 pathway was intact on day 2 (Extended Data Fig. 3C day 2; 64.5 ± 10.1% versus 42.1 ± 6.2%; Extended Data Fig. 2F and G, P = 3.2e-5). This long-term effect of NR-CA1 inhibition was not observed in the control context (Extended Data Fig. 3D day 2; 28.9 ± 14.4% versus 27.3 ± 7.5%). On day 3, freezing in the shocked context in the NR-CA1 inhibited group fell to similar levels as the control context (Shocked: 24.8 ± 11.8% Control: 22.6 ± 9.7%). Freezing levels on day 3 were not significantly different from pre-shock levels in either context (Extended Data Fig. 3C), indicating successful fear extinction. Thus, the absence of the NR-CA1 input on retrieval day 1 caused an increase in CFMR on day 2 in the shocked context, reducing fear extinction, but reinstatement of the NR-CA1 pathway on day 2 allowed extinction to occur on day 3.

We wanted to ensure that the DCZ-induced increase in freezing was not due to a general decrease in movement. To do so, we exposed NR-CA1 inhibited and intact mice to a ‘dark’ context (devoid of any visual cues) for ~5 minutes after they were exposed to both the shocked and control contexts on retrieval days 1-3. In this dark context, mice quickly recovered their running behavior, with NR-CA1 inhibited mice freezing on average only 4.4 ± 4.0% of the time across all 3 days of retrieval. Both within and across-mice controls froze at comparably low levels (on average under 5%; Extended Data Fig. 3B). Therefore, neither DREADD inhibition, nor DCZ itself, impacted the mouse’s ability to move, and the increase in freezing behavior is therefore specific to when mice are navigating in VR contexts. This indicates that the increase in post-shock freezing that we observe in the control context in both NR-CA1 intact and inhibited mice over baseline could not be due to an overall decrease in movement, but is specific to the VR context. It additionally indicates that the increase in fear generalization to the control context in NR-CA1 inhibited mice is due exclusively to NR-CA1 pathway inhibition. Our results indicate that the NR-CA1 pathway sends a potent fear suppression signal, critical for shortening the length of freezing epochs, preventing fear generalization, and inducing contextual fear extinction.

### NR-CA1 Axon Activity Becomes Tuned to Freezing Behavior Following CFC

Excitatory NR projections to the hippocampus are restricted to the stratum lacunosum-moleculare (SLM) of CA1 and subiculum; NR does not project to any other hippocampal subregions or layers^44–46^. Previous work stimulating this projection shows it depolarizes CA1 pyramidal neurons across the dorsal-ventral axis, and induces firing in multiple interneuron subtypes with dendritic processes in SLM^47–50^. However, the activity of the NR-CA1 projection *in vivo* during behavior is unknown. To determine the information conveyed directly from the NR to CA1 during CFMR, we performed *in vivo* 2-photon Ca^2+^ imaging of NR-axons in SLM.

We injected an axon-targeted virus carrying axon-GCaMP6s into NR, followed by a cannula window over CA1 as previously described (Fig. 2A: ^39,40^). Expression in NR was confirmed via histological evaluation following the completion of experiments (Fig 2A; left). NR-axons could be observed in the SLM of CA1 during experiments (Fig. 2A, right) We successfully recorded reliable GCaMP6s expression from 1 NR-axon per mouse (N = 10) in hippocampal CA1 during the VR-CFC paradigm (day 0 and retrieval days 1-3). We limited our analysis to a putative single axon per animal, since all identified axonal segments within the field of view with above-baseline activity were highly correlated (see Method Details). We were additionally able to track a subset of the same NR-axons (N = 4) across days (Extended Data Fig. 5A).

We found that NR axons switched their activity from untuned sparse activity (Fig. 2B; Top) pre-shocks, to activity highly selective for freezing epochs post-shocks, even after filtering for axons with detectable pre-shock activity (Fig. 2B; Bottom; same axon shown on both days). Since behavior necessarily changes following successful CFC, which induces more and longer freezing epochs, we needed to avoid potential confounds in comparing axon activity during dissimilar freezing epoch lengths before and after CFC. To do so, we quantified axon activity in three different ways.

First, we examined the mean normalized *Δf/f* of peaks in both contexts during running (Fig. 2C, Control: Green, Shocked: Blue) and freezing (Fig. 2C, Control: Orange, Shocked: Red). Axons were normalized each day to near-maximum activity, controlling for any potential differences in amplitudes across days. This analysis shows that pre-shocks, NR-axonal activity in the to-be shocked context was similar between running and freezing epochs, with a slight preference for running epochs, which was slightly elevated in the control context (Fig. 2C, Student’s T-test, P = 0.085 in to-be shocked context, 0.039 in control context). Conversely, post-shocks in both contexts and on all 3 retrieval days, we found that NR-axons had significantly higher mean peak activity during freezing compared to running (Fig. 2C, Student’s T-test, Shocked P = day 1: 9.42e-15, day 2: 6.64e-18, day 3, 1.22e-12, Control P = day 1: 6.27e-13, day 2: 7.19e-13, day 3, 1.47e-13).

Second, we binned freezing epochs into 1 s intervals based on their total length from 1-2 s to 6-7 s, and compared pre-shock to post-shocks activity within those bins, therefore comparing the NR-axonal activity of similar lengths of freezing both pre and post shocks (Fig. 2D, Extended Data Fig. 4F). Freezing epochs that were longer in length than 7s were not used for this analysis due to the low quantity of such epochs present pre-shock. (Extended Data Fig. 2C). Within each binned epoch, we trial-aligned activity to the freezing to running transition point (Fig. 2D, black center line). We then compared average NR-axon activity from all axons during freezing (Fig. 2, peach-shaded regions) to activity during running (unshaded regions; Fig. 2D, 3-4 s long freezing epochs shown; all epochs in Extended Data Fig. 4F). Pre-shocks in either context, NR-axons did not significantly modulate their activity between running and freezing epochs (Fig. 2D, Extended Data Fig. 4F). However, post-shocks, we found that NR-axons significantly increased their activity during freezing epochs, compared to reduced activity during running epochs. This was observed during all post-shocks freezing epochs, in both the shocked and control contexts, in all binned intervals (Fig. 2C; Extended Data Fig. 4F).

Third, we characterized the dynamics of NR-axon activity within each freezing epoch on pre-shocks day 0 and compared to retrieval day 1. To do so we aligned NR-axons by dividing each freezing or a running epoch into 5 even bins, each containing a mean normalized *Δf/f* of NR-axon peaks, then took the within-bin mean across all epochs pre and post-shocks. This enabled us to effectively ‘stretch’ or ‘shrink’ all epoch lengths to a uniform standard. Using this method, we found that mean axon activity ramped up rapidly in the beginning of a freezing epoch, plateaued, then fell right before freezing transitioned to running (Fig. 2E). Such temporal dynamics were absent during the freezing epochs pre-shocks (Fig. 2E). These dynamics were all similarly observed in axons tracked across days (Extended Data Fig. 5B-E). These results collectively show that NR-axons projecting to CA1 strongly tune their activity to fearful freezing epochs during CFMR, and this post-shock activity is context-independent.

### Encoding Model Predicts NR-CA1 Axonal Activity, But Only Following CFC

To further quantify the relationship between behavior and NR-axon activity, we developed a quantitative encoding boosted trees decision model to predict axonal activity from behavioral variables. We trained the model using XGBoost^51^ to use behavioral information about freezing epochs, running epochs, velocity, location on the track, and pupil diameter to predict NR-axon activity. We separately trained on 80% of traversals and tested on the remaining 20% of traversals in each mouse, on each day, and in each context (Fig. 3; Extended Data Fig 6; Method Details). Model prediction most heavily relied on behavioral parameters pertaining to whether the mouse was freezing or running, its velocity, and duration passed or remaining within a freezing or running epoch (Fig. 3H, Extended Data Fig. 6B). Overall, the model predicted NR-axon activity well in both the shocked and control contexts on retrieval days post-shocks (with a context/day-combined 0.43 r^2^ goodness of fit; Fig. 3G), but predicted axonal activity poorly in both contexts pre-shocks (with a context-combined 0.01 r^2^; Fig. 3G). In the example mouse shown in panels C-F, the maximum model accuracy pre-shocks was r^2^ of 0.06 (Fig. 3C) compared to a much higher r^2^ of 0.86, 0.88, and 0.78 on retrieval days 1, 2, and 3, respectively (Fig. 3D-F).

Because there was variability in the fluorescence signal recorded from the axons, we checked whether model accuracy was related to the signal-to-noise. Indeed, model accuracy was correlated with axon activity - the greater the change in the normalized fluorescence signal from baseline, the better the model performed (Fig. 3B). The model performed significantly above chance in predicting NR-axon signal in 8/10 mice, on retrieval days 1-3. In 2/10 mice, model prediction was poor on retrieval days, due to lower signal-to-noise ratio (SNR) of the fluorescence signal. However, changes in SNR did not account for the poor model performance pre-shock, as model accuracy was still low in animals with higher axon activity. Although overall activity was higher in post-shock days, pre-shock activity in longitudinally-tracked axons reached similar peak heights as in post-shock days (Extended Data Fig. 4A), and all mice included in analysis had at least 2 peaks reaching a minimum of 0.1 *Δf/f* in the recording session, ensuring that poor model performance was not simply due to a lack of signal to predict. In summary, using an encoding model, we demonstrated that NR-axon activity recorded in hippocampal CA1 can be predicted from freezing behavior during CFMR, but not before the animal is fear-conditioned, revealing the development of predictable structure in NR-axon activity tuned to CFMR.

## DISCUSSION

Our findings expand on a previous canon of work that indicates both the mPFC-NR projection and NR itself are required for contextual fear extinction and preventing fear overgeneralization^29,33–35^. Our results suggest that in addition to these roles, NR reduces time spent freezing following CFC by suppressing CFMR as it is occurring during freezing epochs. We found that the NR-CA1 pathway is a key component of the circuit responsible for mediating the fear suppressive function of NR. This is supported by our observation that NR axons in CA1 become selectively tuned to freezing epochs following CFC and inhibiting the NR-CA1 pathway lengthens freezing epochs. The function of the NR-CA1 pathway in CFMR suppression is not restricted to the context in which shocks were presented, but extends to similar contexts where shocks never occurred. This seems to limit overgeneralization as shown by NR-CA1 inhibition reducing context discrimination. Lastly, the process of suppressing ongoing CFMR by the NR-CA1 pathway also has longer term effects, as shown by reduced fear extinction a day following NR-CA1 inhibition. In summary, our observations support a framework in which the NR-CA1 pathway actively suppresses fear responses by disrupting ongoing hippocampal-dependent CFMR to promote non-fearful behavior, and this process also limits overgeneralization and promotes fear extinction.

Interestingly, we did not observe a significant difference in NR-axon activity during CFMR between contexts on retrieval days, despite NR-CA1 inactivation reducing discrimination between these contexts. It could be that while the NR-axons in CA1 are not contextually modulated, their activity induces postsynaptic dynamics in CA1 that encode differences in context. This is supported by previous work showing that CA1 is specifically necessary for the context-dependence of fear extinction^52^. We also found the difference between NR-axonal activity in freezing and running epochs following CFC does not decrease over days, even as mice decrease their time spent freezing. Previous work shows that extinction does not erase previously-learned contextual fear memories, as reactivation of hippocampal fear memories rapidly reinduces fear behavior^21,53^. This suggests that fear memories are retained but are dormant after extinction. Continued differential activity of NR-axons between freezing and running epochs in CA1, even after extinction, may be necessary to prevent the maladaptive retrieval of dormant fear memories, therefore enabling successful extinction learning.

A recent study showed that during freezing epochs in remote post-conditioning CFMR, optogenetic activation of NR significantly shortened freezing epochs, while inactivation lengthened freezing epochs^54^ - in agreement with our NR-CA1 inhibition results. These authors revealed a transient increase in NR activity before the termination of freezing epochs, and showed a similar signal in the NR-BLA (basolateral amygdala) pathway. The profile of the NR and NR-BLA activity during freezing epochs they report differs from the profile we report, as they ramped up at the end of a freezing epoch and remained high during running. What could be causing this discrepancy? One key difference is the time period in which the NR and NR-BLA signals occur, compared to our reported NR-CA1 signal. We recorded 1 day following CFC, whereas NR and NR-BLA signals were measured 30 days following CFC, a time period in which memories are considered remote and no longer dependent on the hippocampus^55^. In addition, the NR-BLA pathway was not necessary to facilitate extinction one day following shocks. This suggests the NR may interact directly with CA1 to suppress recent CFMR, and directly with the BLA to suppress remote CFMR, through distinct activity dynamics. Future work using closed-loop optogenetic stimulation of the NR-CA1 pathway during freezing epochs, and investigating NR-CA1 activity at remote time points, is needed to directly test this hypothesis.

The input driving the NR-CA1 pathway is most likely from the mPFC, encompassing both the prelimbic (PL) and infralimbic (IL) regions. While PL is needed for fear acquisition and retrieval, IL is necessary for the opposing task of fear suppression and preventing overgeneralization^56–59^. The likely opposing influences of IL and PL on NR during CFMR illustrates the importance of understanding NR output pathways. Our results indicate that a fear suppression signal circuit may be transmitted from IL, through NR, and into CA1 during CFMR. Of note, a small population of NR neurons that project both to CA1 and either PL or IL may have a key role in facilitating cross-regional theta synchrony associated with CFMR^28,60^. While we cannot rule out that some of our recorded NR-axons collaterally project to mPFC, since this population makes up a small subset of all NR neurons (~3-9%^60^), we would expect the majority of our recordings to be from non-dual projecting neurons. It additionally remains to be seen if the NR-CA1 exclusively projecting versus the NR-CA1 dual projecting populations have distinct dynamics during CFMR.

A key question that arises from our work is how the NR-CA1 pathway potentially disrupts CFMR-associated neural dynamics in CA1. NR exclusively projects to the SLM within CA1, where the distal dendritic tuft of pyramidal neurons receive targeted synaptic input from both medial and lateral EC and local inhibitory interneurons^44,61–63^. Whether NR directly synapses on these dendrites is under contention, with contradictory anatomical and electrophysiological reports supporting evidence for and against these direct synapses^29,50,64,65^. Electrophysiological stimulation of NR projections to CA1 in rodent slice work has largely supported that NR projections depolarize, but do not directly drive firing in pyramidal neurons ^48–50,66^, with one notable early exception^47^.

Interestingly, NR and EC both project to the same dendritic compartments, and dual activation of NR and EC projections in slice amplifies nonlinear dendritic spiking, implying that NR/EC interactions may be important *in vivo* for synaptic plasticity^49,67^. Such dendritic-spike-induced plasticity has been associated with the formation of new place fields in novel environments^68^ could provide a mechanism through which NR both disrupts CFMR and promotes extinction learning. Additionally, either NR or EC projections to SLM, when coincident with CA3 inputs through schaffer collaterals, induce burst firing in CA1 pyramidal cells^69–72^. CA1 pyramidal cell bursts are also capable of inducing new place fields in CA1 through behavioral timescale synaptic plasticity (BTSP)^69,71–73^. If our newly-reported NR input to the apical tuft during fearful freezing epochs coincides with CA3 inputs, their combined activity could induce burst firing and initiate BTSP. The bursts themselves could disrupt population dynamics to “push” the network out of CFMR, enabling the behavioral transition from freezing to running, while also inducing new place cell representations to form (remapping) through BTSP to support extinction learning^74^. This framework could explain why inhibiting the NR-CA1 pathway on retrieval day one reduced fear extinction on retrieval day 2. In effect, we may have prevented BTSP from inducing remapping and thus prevented extinction learning.

Alternatively, NR could disrupt CA1 dynamics through inhibition. It is well established that NR induces strong firing in various hippocampal interneuron populations with dendritic processes in SLM^47–50^. Which specific inhibitory populations are directly stimulated by NR is an open question, and one that has wildly divergent implications for the overall impact of NR on CA1 activity. In the case where NR-axons in SLM exclusively target inhibitory interneuron postsynaptic partners, the overall impact of NR-CA1 pathway activation on CA1 pyramidal population activity could still be net inhibitory, net excitatory, or selectively mixed. NR-axons could activate inhibitory micro-circuits that disrupt awake replay of location-specific activity sequences of the shocked context during freezing^75^, or silence temporally-restricted reactivation of engram cells^21^ to induce fear memory suppression and enable extinction learning. Further research on the impact of NR on CA1 dendritic and somatic population dynamics is needed to unravel how the NR-CA1 pathway mechanistically induces suppression of ongoing CFMR.

## SUPPLEMENTAL INFORMATION

## ACKNOWLEDGEMENTS

This work was supported by The Whitehall Foundation, The Searle Scholars Program, The Sloan Foundation, The University of Chicago Institute for Neuroscience start-up funds, a New Innovator grant from the National Institutes of Health (1DP2NS111657-01) awarded to M.S., and a T32 training grant (T32DA043469) from National Institute on Drug Abuse awarded to S.K. We thank the University of Chicago imaging core for assistance with confocal imaging, and the University of Chicago animal care staff for ensuring the well-being of experimental animals. We thank Chad Heer for early help with imaging protocols. We thank Valerie Barreto for helping to train animals and collect confocal data. We thank Timothy Ratigan for assistance with data analysis. We thank Rossten Rad for discussions on experimental design. Finally, we thank Douglas Goodsmith, Jim Heys, Timothy Ratigan, and Rossten Rad for their invaluable comments on previous versions of the manuscript.

## AUTHOR CONTRIBUTIONS

S.K. and M.S. conceived of, designed, and tested the VR-CFC protocol. H.R. modified the VR-CFC protocol in collaboration with S.K. H.R. and M.S. conceived of and designed the experiments. H.R. performed surgeries. H.R. and S.S. collected all *in-vivo* behavioral and imaging data. H.R. and S.S. collected all *post-hoc* data. H.R. wrote the analysis code and analyzed all data. H.R and M.S. interpreted the data and wrote the manuscript, with significant contributions from S.K.

## DECLARATION OF INTEREST

The authors declare no competing interests.

## INCLUSION AND DIVERSITY

One or more of the authors of this paper self-identifies as an underrepresented ethnic minority in science. One or more of the authors of this paper self-identifies as a member of the LGBTQ+ community. While citing references scientifically relevant for this work, we also actively worked to promote gender balance in our reference list. We support inclusive, diverse, and equitable conduct of research.

## KEY RESOURCES TABLE

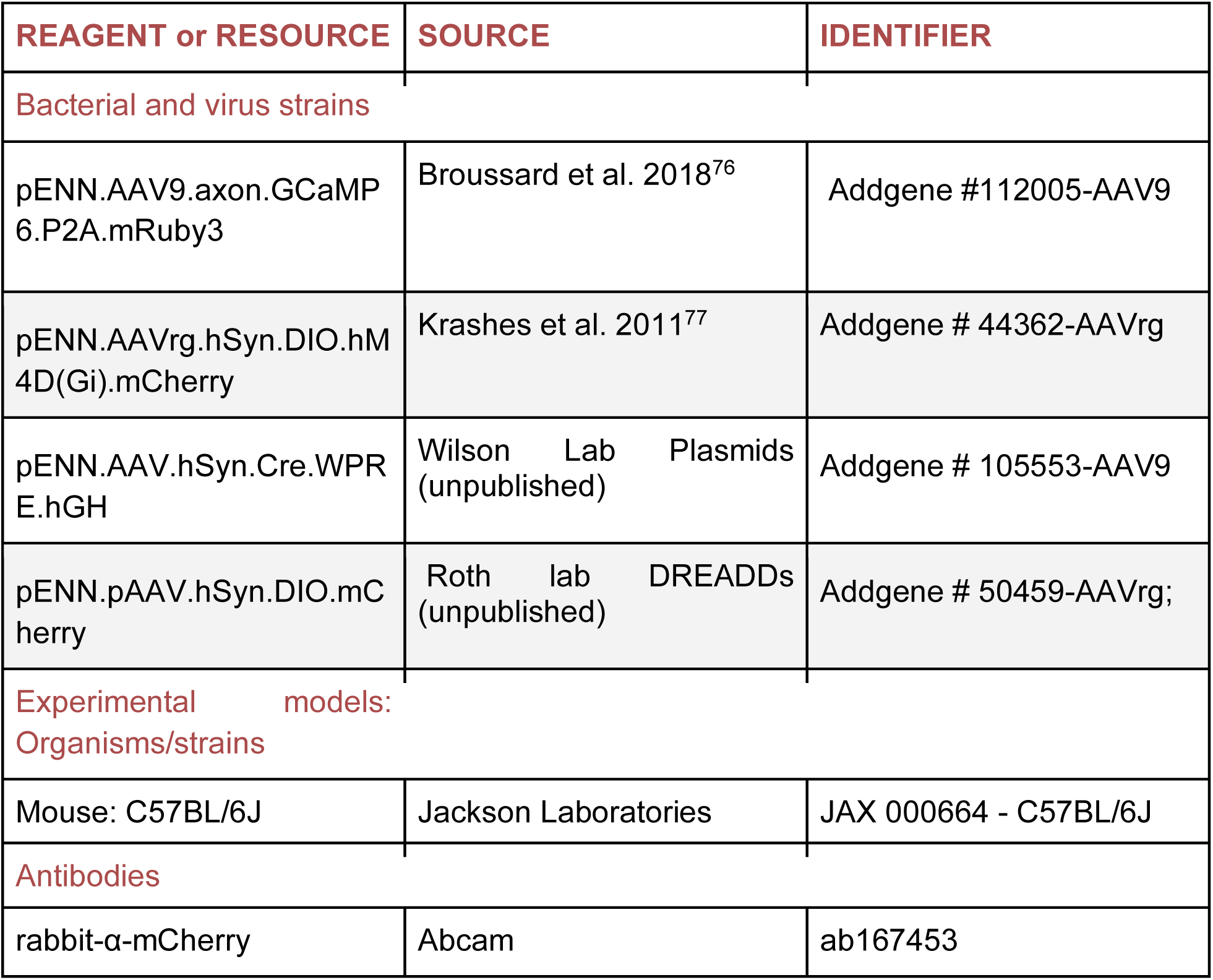
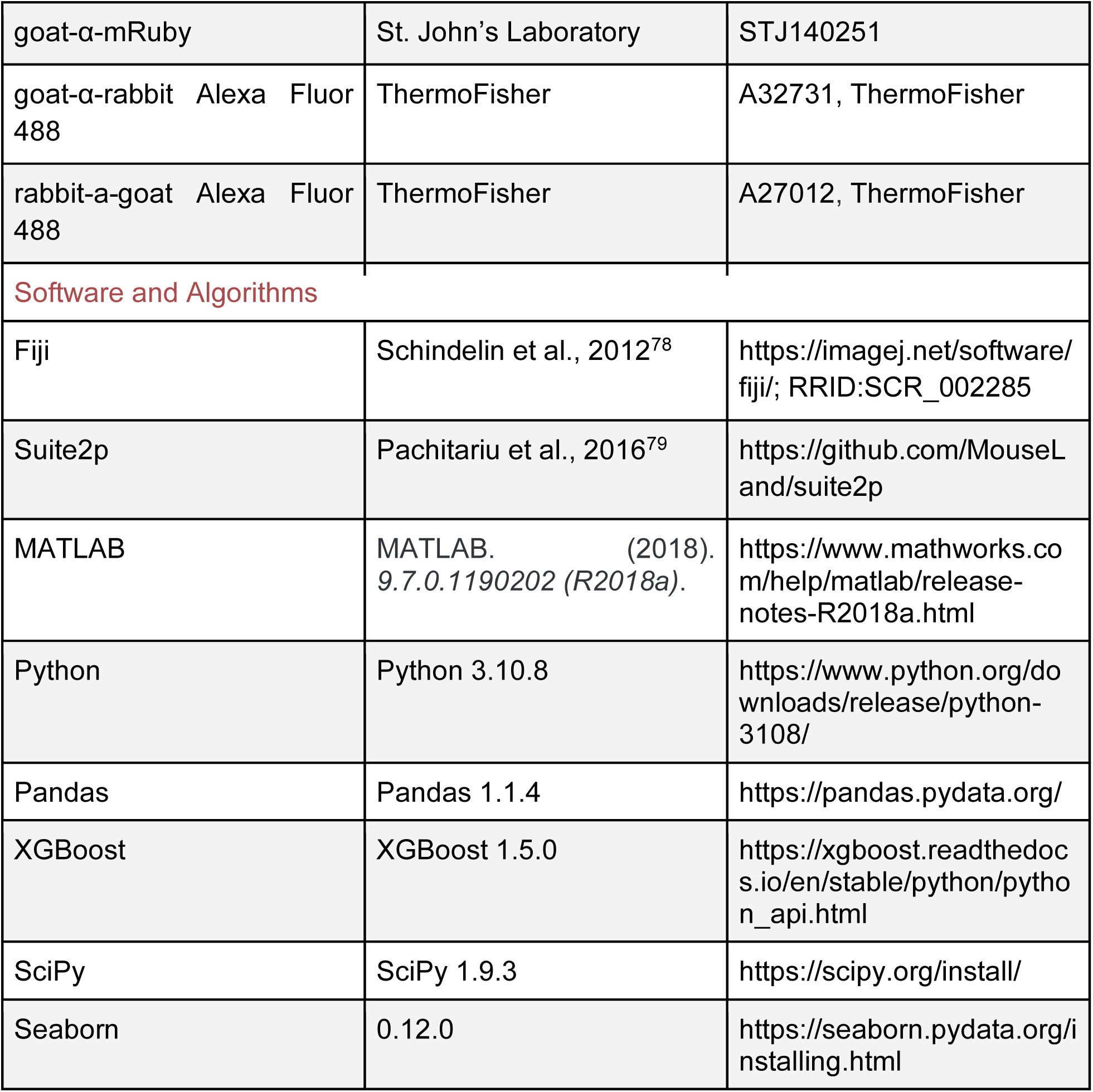

## RESOURCE AVAILABILITY

### Lead contact

Further information and requests for resources and reagents should be directed to the lead contact Mark Sheffield. All unique resources generated in this study are available from the lead contact with a completed Materials Transfer Agreement.

### Materials availability

This study did not generate new reagents.

### Data and code availability

All data reported in this paper and original code will be shared by the lead contact upon request. DOIs are listed in the key resources table. Any additional information required to reanalyze the data reported in this paper is available from the lead contact upon request.

## EXPERIMENTAL MODEL AND SUBJECT DETAILS

All experimental and surgical procedures were in accordance with the University of Chicago Animal Care and Use Committee guidelines. We used 10-20 week old male C57BL/6J wildtype (WT) mice (23-33 g). Male mice were used over female mice due to the size and weight of the headplates (9.1 mm x 31.7 mm, ~2 g) which were difficult to firmly attach on smaller female skulls, and low weights reached under water restriction in female mice making the additional weight of the tailcoat potentially burdensome and interfere with experimental results. Mice were individually housed in a reverse 12 hour light/dark cycle and behavioral experiments were conducted during the animal’s dark cycle. We are unaware of any influence of strain or sex on the parameters analyzed in this study. A total of 66 mice were used, 36 of which were used in the final data. 30 mice did not meet running criteria (See Method Details: Behavior and Virtual Reality). Of these 8 never reached the 4 traversals/minute cutoff after 14+ days of training, and 22 did not meet movement criterion after removal of water reward and addition of tailcoat. 20 mice were used in the NR-CA1 intact group, 10 of which were imaged in the NR-axon NR-CA1 intact group. In the remaining 16 mice, 4 were eliminated for z-motion drift. The remaining 12 mice were used for control groups: 4 mice were used for no-shock control, 4 mice were used for mCherry DREADD control, and 4 mice were used for saline DREADD control.

## METHOD DETAILS

### Mouse surgery and viral injections

Mice were anesthetized (~1-2% isoflurane) and injected with 0.5 ml of saline (intraperitoneal IP injection) and 0.5 ml of Meloxicam (1-2 mg/kg, subcutaneous injection) before being weighed and mounted onto a stereotaxic surgical station (David Kopf Instruments). A small craniotomy (1-1.5 mm diameter) was made over the hippocampus (± 1.7 mm lateral, −2.3 mm caudal of Bregma) or nucleus reuniens (0.0 lateral, −0.6 caudal of Bregma). For NR imaging experiments, an axon targeted genetically-encoded calcium indicator, AAV9-axon-GCaMP6s-P2A-mRuby3 (pAAV-hSynapsin1-axon-GCaMP6s-P2A-mRuby3 was a gift from Lin Tian Addgene viral prep # 112005-AAV9; http://n2t.net/addgene:112005; RRID:Addgene_112005) was injected (~50 nL at a depth of 4.1 mm below the surface of the dura) using a beveled glass micropipette leading to GCaMP6s and mRuby expression in a population of NR neurons. For DREADD experiments, first AAVrg-hSyn-DIO-hM4D(Gi)-mCherry (pAAV-hSyn-DIO-hM4D(Gi)-mCherry was a gift from Bryan Roth Addgene viral prep # 44362-AAVrg; RRID:Addgene_44362) was injected into bilateral hippocampal CA1 SLM (~50 nL per side at a depth of −1.5 mm below the surface of the dura). In the same surgical procedure, AAV9-hSyn-Cre (pENN.AAV.hSyn.Cre.WPRE.hGH was a gift from James M. Wilson Addgene viral prep # 105553-AAV9; http://n2t.net/addgene:105553; RRID:Addgene_105553) was injected into bilateral NR (~100 nL at a depth of −4.1 mm). For NR-DREADD Controls, AAVrg-hSyn-DIO-mCherry (pAAV-hSyn-DIO-mCherry was a gift from Bryan Roth Addgene viral prep # 50459-AAVrg; http://n2t.net/addgene:50459; RRID:Addgene_50459) was substituted for AAVrg-hSyn-DIO-hM4D(Gi)-mCherry. Afterwards, the site was covered using dental cement (Metabond, Parkell Corporation) and a metal head-plate (9.1 mm x 31.7 mm, Atlas Tool and Die Works) was also attached to the skull with the cement. Mice were separated into individual cages and water restriction began the following day (0.8-1.0 ml per day). At least 7 days following injection surgery, and approximately 7 days prior to the beginning of mouse training, mice underwent another surgery to implant a hippocampal window as previously described^80^. Following implantation, the head-plate was reattached with the addition of a head-ring cemented on top of the head-plate which was used to house the microscope objective and block out ambient light. Post-surgery mice were given 1-2 ml of water/day for 3 days to enhance recovery before returning to the reduced water schedule (0.8-1.0 ml/day). Expression of axon-GCaMP6s reached a steady state ~50 days after the virus was injected, as monitored through 2p imaging. Expression of hM4D(Gi)-mCherry was validated using *post-hoc* confocal imaging.

## Behavior and Virtual Reality

Our virtual reality (VR) and treadmill setup was designed similarly to previously described setups^40^. The virtual environments that the mice navigated through were created using VIRMEn^81^. Mice were head restrained with their limbs comfortably resting on a freely rotating styrofoam wheel (‘treadmill’). Movement of the wheel caused movement in VR by using a rotary encoder to detect treadmill rotations and feed this information into our VR computer, as in (Heys et al., 2014; Sheffield et al., 2017). During training, mice received a water reward (4 µl) through a waterspout upon completing each traversal of the track (a lap), which was then associated with a clicking sound from the solenoid. Upon receiving the water reward, a short VR pause of 1.5 s was implemented to allow for water consumption and to help distinguish traversals from one another rather than them being continuous. Mice were then virtually teleported back to the beginning of the track and could begin a new traversal. Mice were also teleported to the beginning of a new contextual exposure.

Mouse behaviors (running velocity, track position) were collected using a PicoScope Oscilloscope (PICO4824, Pico Technology). Pupil tracking was done through the imaging software (Scanbox, Neurolabware) at 15.49 Hz, using Allied Vision Mako U-130b camera with a 25 mm lens and a 750 nm longpass IR filter. IR illumination from the objective was used to illuminate the pupil for tracking. Behavioral training to navigate the virtual environment began ~7 days after window implantation (~30 minutes per day) and continued until mice reached a speed of greater than 4 traversals per minute, which took 10-14 days (although some mice never reached this level). This high level of training was necessary to ensure mice continued to traverse the track similarly after reward was removed. Initial experiments showed that mice that failed to reach this criterion typically would not traverse the track as consistently without reward^40^ a potential confound for post-shocks freezing data (data not shown). Mice that did not reach this criterion were not used for these experiments (28 mice removed across all conditions).

### Contextual Fear Conditioning Paradigm

In mice that reached criteria in the training environment (>4 traversals per minute), were first exposed to two novel environments without water reward for 322 s (~5 minutes) each, with the addition of a custom-made tailcoat made of conductive fabric (Adafruit). Only mice that continued to maintain a speed of 4 traversals > minute without water rewards and with the tailcoat on were allowed to continue the experiment. Subselecting for mice with this consistent running behavior helped us to ensure that freezing responses recorded later were not due to the presence of the tailcoat or any discomfort from head-fixation or removal of reward. Here onwards, the tailcoat was kept on the mouse during the experimental sessions on all subsequent experimental days. Each contextual exposure was for a duration of ~ 5 minutes. Prior to experimental day 0, mA level of shock delivery was confirmed using an oscilloscope. On day 0, mice were exposed to both novel contexts, then shocked in one of the two contexts, administering 6 0.6mA shocks delivered at an interval of 20-26 seconds each, (Coulbourn Instruments Precision Animal Shocker). Mice displayed rapid sprinting behavior when they received the tail shock, allowing us to confirm the delivery of shocks in real-time (Extended Data Fig. 1D). On subsequent days, mice were exposed to both the shocked and non-shocked (control) contexts pseudorandomly, for 3 days.

### DREADD Experimental Protocol

To activate the hM4D(Gi) receptor and silence a subset of NR glutamatergic neurons that project to CA1, we used Deschloroclozapine dihydrochloride (DCZ, MedChemExpress). Due to the slow kinetics and known off-target effects of CNO and high, rapid efficacy of DCZ^43^, we chose to use DCZ for inactivation as in our past work^40^. Once mice met training criteria, they were habituated to the injection process. They were exposed to the rewarded training environment for ~10 min. Afterwards, they were removed from the VR set up, placed in the holding room, and injected with ~150 µL of a 12% DMSO/Saline solution. After ~30-45 min, they were placed back in the VR setup and exposed to the rewarded training environment again for an additional 10 min. This was repeated for 3 days to acclimate mice to the injection procedure. Mice additionally received ~150 µL of a 12% DMSO/Saline solution on Day −1 of the experiment 30 minutes prior to first exposure to both neutral contexts to mimic conditions on Day 0.

For animals receiving DCZ injections, i.e. both the experimental NR-CA1 inhibited AAVrg-hSyn-DIO-hM4D(Gi)-mCherry group and the control NR-CA1 intact AAVrg-hSyn-DIO-mCherry group, DCZ was dissolved in DMSO at at .02 mg/mL concentration and stored at −80 °C on day 0. On retrieval day 1, DCZ solutions were thawed to room temperature and diluted to 0.01 mg/mL with DMSO/Saline. ~30 minutes prior to context exposure, mice were brought to a holding room and IP injected with 0.1 mg/kg DCZ of a .02 mg/mL solution. A separate control group with hM4Di expression intact received DMSO/saline instead of DCZ on retrieval day 1. These mice were injected with a weight matched quantity (~100-150 µL) of saline in place of 0.01 mg/mL DCZ. In all groups, a quantity of DMSO/Saline solution identical to IP injection amount on Day 1 (~100-150 µL) was injected on all other experimental days, ~30 minutes prior to imaging, to control for the impact of any potential IP injection-induced stress. Imaging protocol for all DREADD NR-CA1 inhibited experimental mice and intact controls was kept identical to VR-CFC NR-axon imaged mice, with the addition of a ‘dark’ imaging session after context exposures of the same duration, where no context was displayed on screens, for ~5 minutes, to check for any impact of DCZ on movement (Extended Data Fig. 3B).

### Two-photon imaging

Imaging was done using a laser scanning two-photon microscope (Neurolabware). Using a 8 kHz resonant scanner, images were collected at a frame rate of 15.49 Hz with unidirectional scanning through a 16x/0.8 NA/3 mm WD water immersion objective (MRP07220, Nikon). axon-GCaMP6s was excited at 920 nm and mRuby was excited at 1040 nm with a femtosecond-pulsed two photon laser (Insight DS+Dual, Spectra-Physics) and emitted fluorescence was collected using two GaAsP PMTs (H11706, Hamamatsu). The average power of the laser measured after the objective ranged between 60-100 mW, and was kept constant across days of imaging. A single imaging field of view (FOV) was positioned between 350-500 µm below the putative surface and 400-700 µm equally in the *x/y* direction to collect data from as many NR axonal segments as possible. Time-series images were collected through Scanbox (Neurolabware) and the PicoScope Oscilloscope was used to synchronize frame acquisition timing with behavior. When possible, the same axonal field was returned to across days (Extended Data Fig. 5, N = 4/10 imaged mice).

### Immunohistochemistry and Confocal Imaging

Expression of either hm4D(Gi)-mCherry or GCaMP6s-mRuby in glutamatergic neurons in NR were checked *post hoc*. Mice were anesthetized with isoflurane and perfused with ~10 ml phosphate-buffered saline (PBS) followed by ~20 ml 4% paraformaldehyde in PBS. Brains were removed and immersed in 30% sucrose solution overnight before being sectioned at 30 µm-thickness on a cryostat. Brain slices were collected into well plates containing PBS. Slices were washed 5 times with PBS for 5 min then were blocked in 1% Bovine Serum Albumin, 10% Normal goat serum, 0.1% Triton X-100 for 2hrs. Brain slices were then incubated with either 1:500 rabbit-α-mCherry (ab167453, Abcam) or 1:500 goat-α-mRuby (STJ140251, St John’s Laboratory) in a blocking solution at 4°C. After 48 hrs, the slices were incubated with either 1:1000 goat-α-rabbit Alexa Fluor 488 secondary antibody (A32731, ThermoFisher) or 1:1000 rabbit-a-goat Alexa Fluor 488 secondary antibody (A27012, ThermoFisher) respectively, for 2 hrs. Brain slices were then collected on glass slides and mounted with a mounting media with DAPI (SouthernBiotech DAPI-Fluoromount-G Clear Mounting Media, 010020). Whole-brain slices were imaged under ×10 and x40 with a Caliber I.D. RS-G4 Large Format Laser Scanning Confocal microscope from the Integrated Light Microscopy Core at the University of Chicago.

## QUANTIFICATION AND STATISTICAL ANALYSIS

### Image Processing and ROI selection

Time-series images were preprocessed using Suite2p (Pachitariu et al., 2017). Movement artifacts were removed using rigid and non-rigid transformations and assessed to ensure absence of drifts in the *z*-direction. Datasets with visible *z*-drift were discarded (N = 4). All datasets collected during shock administration on Day 0 were discarded, due to the high velocity post-shocks sprinting behavior of mice making FOVs too unstable for reliable analysis. Regions of interest (ROIs) were also defined using Suite2p (Fig. 1Aiii) and manually inspected for accuracy. Baseline corrected *Δf/f* traces across time were then generated for each ROI.

In addition, to control for in-experiment motion artifacts for small axonal segments, a red mRuby channel was recorded simultaneously to GCaMP6s channel recordings. Per ROI, a savitzky-golay filter was applied to both channels to smooth the signal. Then, the demeaned red channel was ‘subtracted’ from the demeaned green channel, by orthogonalizing their vectors in variance space. That is, we took the projection of the red channel onto the green channel as 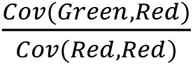. *Red*, and then subtracted that vector from the green channel. This results in a new vector which is guaranteed to have zero covariance with the red channel, thus removing any linear effects of the background fluorescence on the trace. All ROIs were analyzed for covariance, and any ROIs exceeding the 99th percentile of a shuffle distribution were combined using PCA and the first PC taken, in a method similar to ^82^. To ensure traces had sufficient activity for analysis, all mice used were required to have one axon per FOV with activity that exceeded 10% *Δf/f* twice on each experimental day (N = 10 mice). The activity of each axon was then internally rescaled per day to the 99th percentile of max activity to account for inter-axonal differences in calcium brightness. Peaks were calculated using the scipy.signal.find_peaks package with a required minimum height of 10% *Δf/f,* distance of 0.5 s, and prominence of 0.1. Multiple segments per mouse were not used, as correlation remained high enough in mice with multiple differentiable segments (>0.2) to not rule out that these segments could have originated from the same original axonal projection.

### Pupil measures

To obtain images with dark pupils and high contrast around the borders of the pupils, pupil images were inverted, and their brightness/contrast was adjusted in ImageJ. Pupil area, pupil center of mass (COM), Pupil x and y positions, and blinking area were obtained using FaceMap (Stringer et al. 2019). Pupil data during blinking periods (frames where blinking area < mean – twice the standard deviation of the blinking area) was removed and the pupil data was interpolated to match the 2-photon imaging frame rate (15.49 Hz). Pupil area and x and y position data were smoothed with a savitzky-golay filter.

### Boosted Trees Model

The encoding model used is the python implementation of the open-source gradient boosted trees algorithm XGBoost^51^. Behavioral model parameters (described below) were used to predict axon trace values. For reproducibility, the seed was set to 42. Data were then split into laps, and split using an 80/20 train/test regime. Model was run either per mouse (Fig. 3) or across mice (Extended Data Fig. 6C), per, day, and context paradigm, for a total of 8,000 runs (N=10 mice, 4 days, 2 contexts, 100 draws). Chance performance was determined by shuffling neural activity by traversal compared to behavioral readout per mouse, across contexts and days. Model hyperparameters were set to: gamma = 1, learning_rate=0.01, n_estimators = 1000, base_score = 1, early_stopping_rounds = 5. The coefficient of determination *r^2^* is defined as 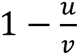 where *u* is the residual sum of squares Σ (*y_true_* - *y_prod_*)^2^ and *v* is the total sum of squares Σ (*y_true_* - *y_mean_*)^2^. The best possible *r^2^* score is 1.0, and the *r^2^* score can be negative because the model can be arbitrarily worse than chance. For ease of interpretability in Fig 3h, the following groupings of related behavioral variables were made and their contributions averaged within each group: freezing = (‘freeze’, ‘is freezing’, ‘freeze remaining’, ‘is postfreeze’, ‘freeze progress’, ‘freeze elapsed’), velocities = (‘recorded velocity’, ‘velocity back 15 frames’, ‘velocity back 8 frames’, ‘velocity forward 8 frames’, ‘velocity forward 15 frames’), running = (‘is running’, ‘running progress’,’running remaining’, ‘is backtracking’,’running_elapsed’, interval = (‘interval_elapsed’, ‘interval_remaining’,’interval_progress’, location = ‘location’, pupil = (‘pupil area’, ‘pupil x position’, ‘pupil y position’). We used the importance type ‘gain’ parameter to determine the importance of each figure to the model’s overall performance. ‘Gain’ is how much an individual feature contributed to model accuracy (i.e. the distance between predicted and actual r^2^ values) on each branch. For each feature’s use in the model, that value is summed, then averaged across all models by context. Full gain fractions for each parameter are shown in Extended Data Fig. 6B.

### Behavioral Parameters and Quantifications

All parameters described below were calculated per mouse, day, and context, and used in model training, with the exceptions of total displacement and shocks.

#### Time to complete a traversal

This was calculated as the total time (in seconds) taken by the animal to run from 0 to 200 cm. Frames recorded within the teleportation window were dropped from analysis.

#### Total Displacement

Total displacement was calculated as the distance traversed per mouse, per context, per day.

#### Freezing

Freezing epochs were determined as uninterrupted epochs where mouse velocity fell below 0.001 cm/s for at least 12 consecutive frames (~0.75 s). All epochs of velocity below 0.001 cm/s but not reaching 12 consecutive frames were not considered freezing or running, and were discarded from future analysis. Freezing epochs were then counted up, and each not in a freezing epoch assigned a ‘0’, while each frame in a freezing epoch given a numeric value corresponding to the number of epochs in that recording (i.e. all frames that contained the 4th freeze of the recording would be assigned the integer ‘4’). Subsequent freeze features were then calculated, including the binary variable ‘is freezing’ which assigns a 1 to frames considered freezing, and 0 to frames not considered freezing, two sawtooth functions ‘freeze remaining’, and ‘freeze elapsed, which counts the frames from the beginning of a freeze up or down until the end of a freeze, respectively, and ‘freeze progress’ which tracks the progress of a freeze as a fraction from 0 to 1.

#### Running

Running was determined as any epoch where forward progress velocity was sustained over 0.001 cm/s for 2 consecutive frames. The variables ‘is running’, ‘running remaining’, ‘running elapsed’, and ‘running progress’ are calculated using the running epoch data in the same fashion as their freezing counterparts.

#### Backward movement

Some mice demonstrated backward movement behavior in the virtual environment post-shocks, where they made backwards movement through the context. This behavior was analyzed separately from running or pausing in Extended Data Fig. 1E. The binary variable ‘is backtracking’ assigns a 1 to frames considered backtracking, and 0 to frames not considered backtracking.

#### Shocks

Shock delivery was recorded through the Picoscope. Shock location on track and stereotyped post-shocks sprinting behaviors are quantified in Extended Data Figure 1B and 1D.

#### Velocity

Velocity was both directly measured through the picoscope encoder, and recalculated from position, to assess for accuracy. Recorded velocity was used for all velocity calculations and model training. Values were converted into cm/s for presentation.

#### Velocity offsets

Future and past velocity at ~1 s and ~0.5 s were calculated by offsetting the velocity to frames. The resulting non-existent 8 or 15 velocity frames at the beginning or end of the trace were extrapolated from the prior 15 frames.

#### Acceleration

Acceleration was calculated as the first derivative of recorded velocity.

#### Intervals

Three variables, ‘interval elapsed’, ‘interval progress’, and ‘interval remaining combine pausing and running information into one datastream. Interval elapsed takes the component parts ‘freeze elapsed’ and ‘running elapsed’, and counts the time elapsed in either a pausing or running interval, before resetting at a switch point. ‘Interval remaining’ and interval progress do the same calculation, but using freeze remaining’/’running remaining’ and ‘freeze progress’/’running progress’

#### Location

Animal’s position on virtual track was determined for each frame, and binned in 1 cm bins along the virtual track.

#### Pupil Area

Pupil area was calculated by FaceMap as previously described, then filtered with a savitzky-golay filter for smoothing.

#### Pupil horizontal (x) movement

Pupil x movement was calculated by FaceMap as previously described then filtered with a savitzky-golay filter for smoothing.

#### Pupil vertical (y) movement

Pupil y movement was calculated by FaceMap as previously described, then filtered with a savitzky-golay filter for smoothing.

### Statistics

For data distributions, a Shapiro-Wilk test was performed to verify if the data was normally distributed. For non-normal distributions, a paired Wilcoxon signed rank test, unpaired Student’s T test, or an unpaired Mann-Whitney U test was used. For samples with five data points or less, only a non-parametric test was used. Multiple comparisons were corrected with a post-hoc holm-sidak correction. Box and whisker plots were used to display data distributions where applicable. The box in the box and whisker plots represent the first quartile (25^th^ percentile) to the third quartile (75^th^ percentile) of the distribution, showing the interquartile range (IQR) of the distribution. The black line across the box is the median (50^th^ percentile) of the data distribution. The whiskers extend to 1.5*IQR on either side of the box. A data point was considered an outlier if it was outside the whiskers or 1.5*IQR. Significance tests were performed with and without outliers. Data distributions were considered statistically significant only if they passed significance (p < 0.05) both with and without outliers. Significance numbers reported are without outliers. To model the probability distribution in the datasets and get an accurate idea of the data shape, a kernel density estimate was fitted to the data distribution and is shown alongside histograms. Cumulative probability distribution functions were compared using a Kolmogrov-Smirnov test. Correlations were performed using Pearson’s correlation coefficient. p < 0.05 was chosen to indicate statistical significance and p-values presented in figures are as follows: ∗, p < 0.05, ∗∗, p < 0.01, ∗∗∗, p < 0.001, N.S. not significant. Darker lines in the center of line plots are the mean, and shading is the 95% confidence interval, unless stated otherwise in text or figure legends. All regression analysis was conducted using the statsmodels Robust Linear Model package, which estimates a robust linear model via iteratively reweighted least squares, given a robust criterion estimator. The M-estimator minimizes the function 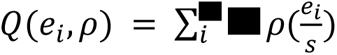 where ρ is a symmetric function of the residuals and *s* is an estimate of scale. We used standardized median absolute deviation for *s* and Huber’s loss function, as it is less sensitive to outliers. Shading on regressions indicate 95% CI. (see https://www.statsmodels.org/dev/examples/index.html#robust-regression for additional details). Data preprocessing was done with MATLAB (Mathworks, Version R2018a). All other data and statistical analyses were conducted in Python 3.7.4, with primary data accrued in Pandas DataFrames, and data figures were made in Python 3.7.4 using the Seaborn and Matplotlib packages (https://www.python.org/). Schematic figures (Fig 1a, Fig 1b. Fig 2a, Fig 3a, and Extended Data Fig 1a), some figure text, and figure layouts were made with BioRender (https://biorender.com/).

---

**Extended Data Figure 1.**
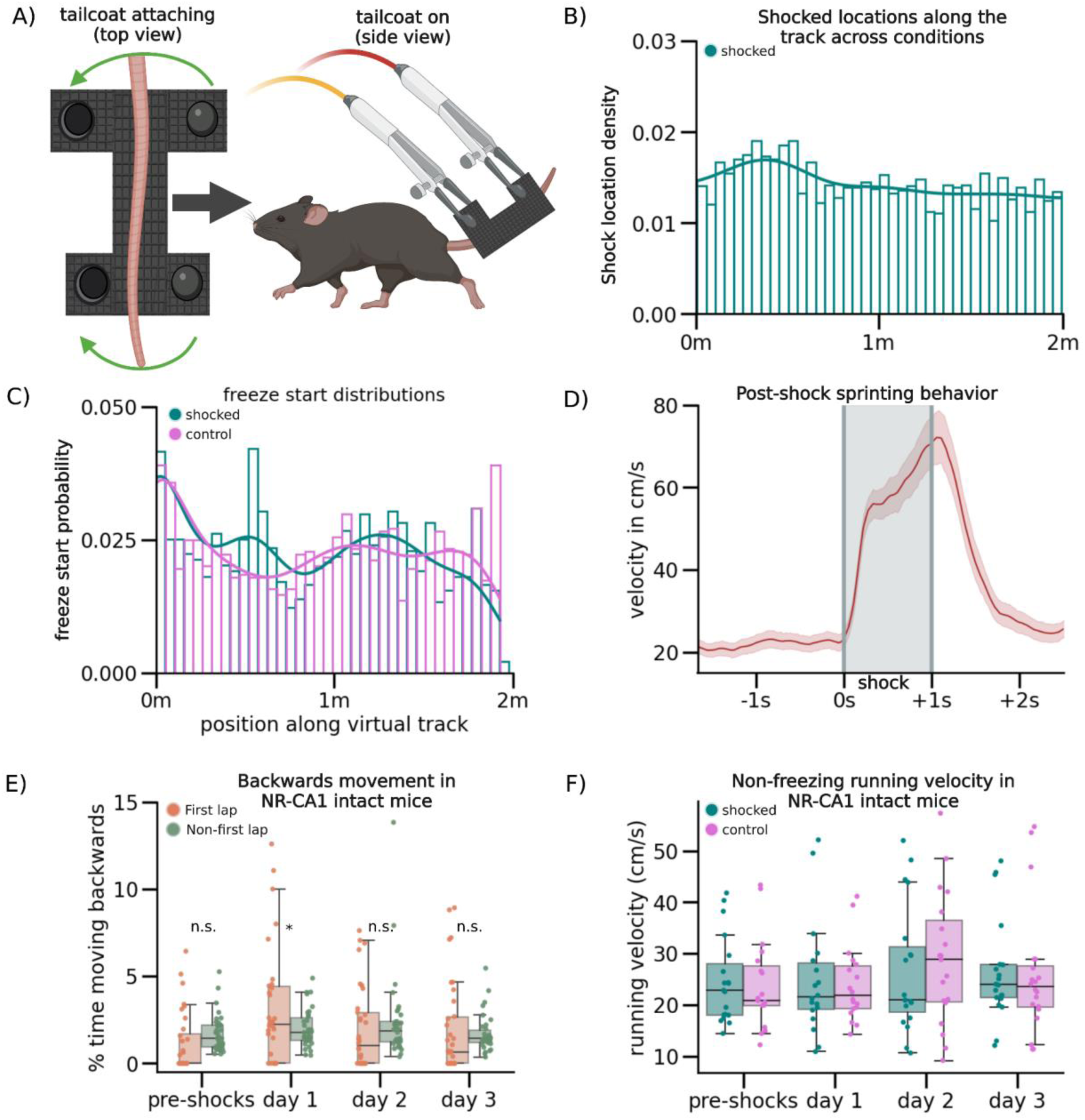
Shock delivery and behavioral responses associated with VR-CFC. (A) Schematic of tailcoat apparatus. Left: Tailcoat is custom-made out of conductive cloth (Adafruit) with two metal snaps that secure the cloth around the tail. Right: The tail coat is wrapped around the mouse’s tail and suspended using lightweight alligator clips forming a supportive ‘hammock’ structure, which are connected to a device that generates electric shocks. In this way, the circuit is completed via the portion of the tail wrapped by the conductive fibers which is where the animal will experience the shock. This entire apparatus weighs less than 1.8g the weight of which is supported by the ‘hammock’ structure, ensuring minimum disruption of the mouse’s running behavior following the addition of the tail cloth. B) Density histogram of shock locations across the virtual track. Track length was split into 40 bins, and locations where shocks were administered were registered as 1 throughout the duration of the shock administration, while all other locations were registered as 0, in all mice that were shocked across conditions. Y-axis is the probability distribution of shocked locations containing a freeze start. Mice were shocked evenly at pseudorandom locations across the context, as no section of track received significantly more shocks than any other section of track (Kolmogorov-Smirnov, P =0.83). (C) Density histogram of freeze start locations across the virtual track. Track length was split into 40 bins and mouse freezing start locations across conditions and post-shocks days were binned with their probability of occurrence plotted. These data indicated that mice froze evenly across the track and indistinguishably across contexts (Kolmogorov-Smirnov, P=Shocked: 0.056, Control: 6.13e-1, Mann-Whitney U, P=0.80). (D) Average post-shocks velocity shows post-shocks sprinting behavior (line = mean, red shading = 95% CI). Grey shading indicates duration of shock. Mouse velocity was aligned to shock initiation and plotted across all shocks in all shocked conditions. When mice received a shock, they briefly ‘sprinted’, nearly quadrupling their velocity for the duration of shocks and briefly post-shocks, before returning to baseline. We hypothesize this sprinting behavior is a stereotyped escape behavior from receiving a shock on the tail. Because this sprinting behavior perfectly correlated with shock onset, we also used it as a real-time verification for whether or not a mouse received the shock. (E) Backward movement behavior in NR-CA1 intact mice. Dots indicate average percent time spent moving backwards in each epoch, either in the first traversal (green), or all other traversals (orange), boxplot indicates median, 25-75th interquartile range, whiskers include all data points not determined to be outliers. We observed instances of backwards movement behavior, where mice attempted to move ‘backwards’ on the track. This was significantly more common in the first 1-3 traversals of the track than all other traversals post-shocks (Student’s T, P=pre-shocks: 0.51, day 1: 0.0023, day 2: 0.30, day 3: 0.01), more common post-shocks than pre-shocks (Student’s T, P=pre-shocks: 0.31, post-shocks: 0.0039), and more common in the feared context than the control context on day 1 (Student’s T, P=0.004). We interpret this backwards movement as an attempt by the mouse to exit the context by ‘backing out’ of it, and classify it as a fearful behavior. (F) Dots indicate each mouse’s mean non-freezing velocity per context, boxplot indicates median, 25-75th interquartile range, whiskers include all data points not determined to be outliers. We calculated mean running velocity in each mouse across contexts, conditions, and days in identified running epochs to determine if our VR-CFC protocol altered running speed when the mice were not freezing. We did not observe any significant differences in velocity between contexts or conditions across days.

**Extended Data Figure 2:**
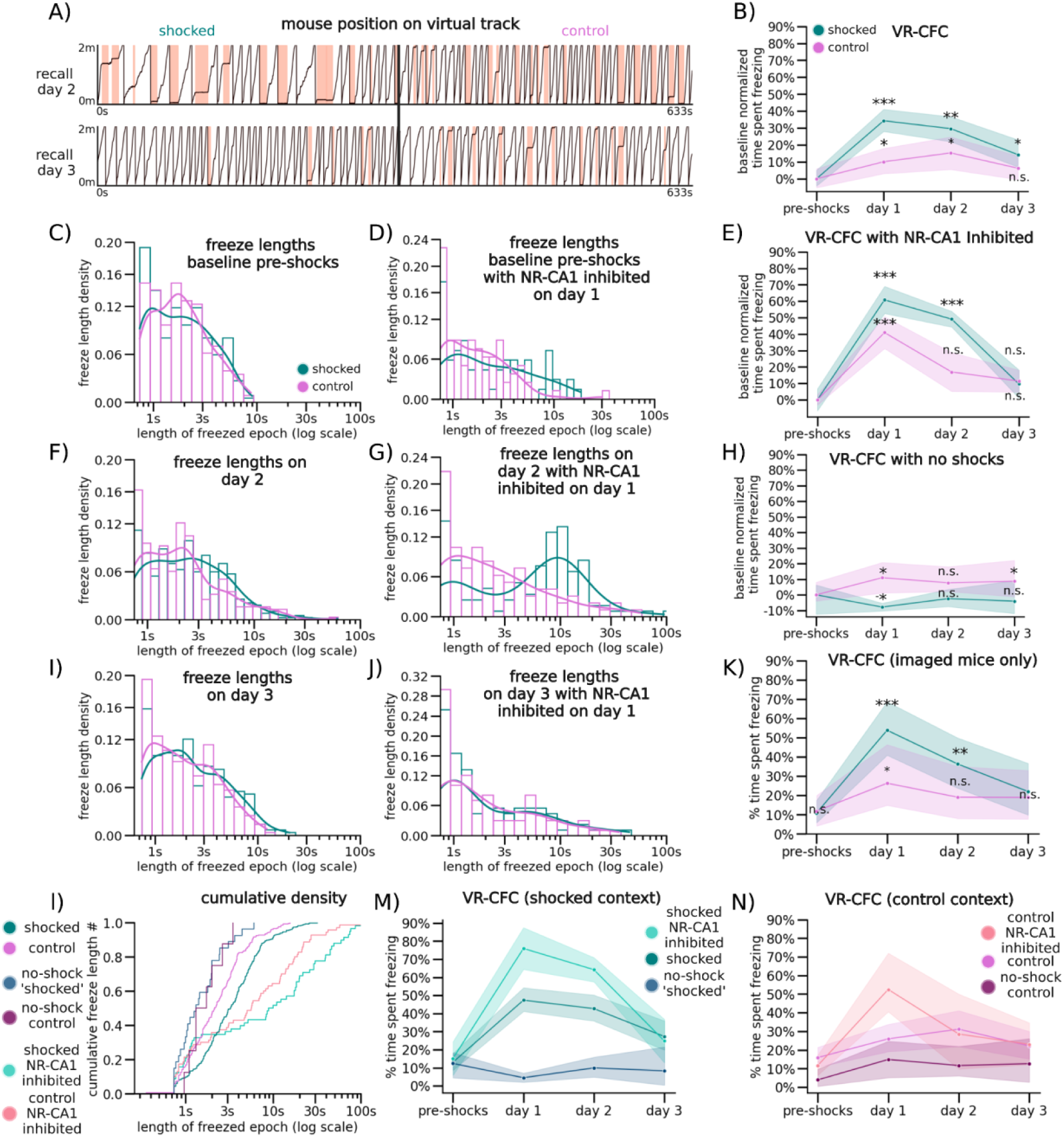
Additional VR-CFC behavioral analysis in shocked and control contexts with and without NR-CA1 pathway inhibition. (A) Example traces from retrieval day 2 (Top) and retrieval day 3 (Bottom) from the same mouse and displayed in the same fashion as Fig. 1C. (B) Instead of comparing between contexts, we normalized mouse percent time freezing to baseline in each context, and tested the difference from baseline within context (N = same 20 mice as Fig. 1; CI = 95% shaded area). In this comparison, shocked mice froze significantly more in the shocked context post-shocks than pre-shocks on both retrieval days 1 and 2. By retrieval day 3, freezing in the feared context remained slightly elevated, while freezing in the control context returned to baseline (Wilcoxon Rank Sum was performed comparing percent time spent freezing per day within each context to the pre-shock baseline with holm-sidak multiple comparisons corrections, P = Shocked: 5.04e-07, 3.89e-07, 4.92e-02, Control: 3.23e-02, 1.92e-02, 3.27e-01). (C) Freeze lengths for the pre-shock baseline day for NR-CA1 intact mice, calculated as in Fig. 1G. (D) Freeze lengths for the pre-shock baseline day for NR-CA1 DREADD inhibited mice, calculated as in Fig. 1H. (E) Equivalent of Fig. 1E with baseline normalized comparisons calculated as in Sup. Fig 2b. (F) Freeze lengths for retrieval day 2 for NR-CA1 intact mice, calculated as in Fig. 1G. (G) Freeze lengths for retrieval day 2 for NR-CA1 DREADD inhibited mice, calculated as in Fig. 1H. (H) Equivalent of Fig. 1F with baseline-normalized comparisons calculated as in Sup. Fig 2b. (I) Freeze lengths for retrieval day 3 for NR-CA1 intact mice, calculated as in Fig. 1G. (J) Freeze lengths for retrieval day 3 for NR-CA1 DREADD inhibited mice, calculated as in Fig. 1H. (K) Equivalent to Fig. 1D, but only with the subset of mice that were imaged from (N=10 mice), comparisons calculated as in Fig. 1D. (L) Cumulative density plot of freeze lengths in both contexts and three conditions. (M) Percent time freezing in only the shocked context across three conditions, NR-CA1 inhibited (Top; bright teal), NR-CA1 intact (Middle; teal), and NR-CA1 intact in mice that did not receive shocks in any context (Bottom; navy). These lines are the same data from Fig 1D-F, replotted together for effective visualization of condition on freezing behavior in the shocked context. (N) Percent time freezing in only the control context across three conditions, NR-CA1 inhibited (Top; apricot), NR-CA1 intact (Middle; pink), and NR-CA1 intact in mice that did not receive shocks in any context (Bottom; burgundy). These lines are the same data from Fig 1D-F, replotted together for effective visualization of condition on freezing behavior in the control context.

**Extended Data Figure 3:**
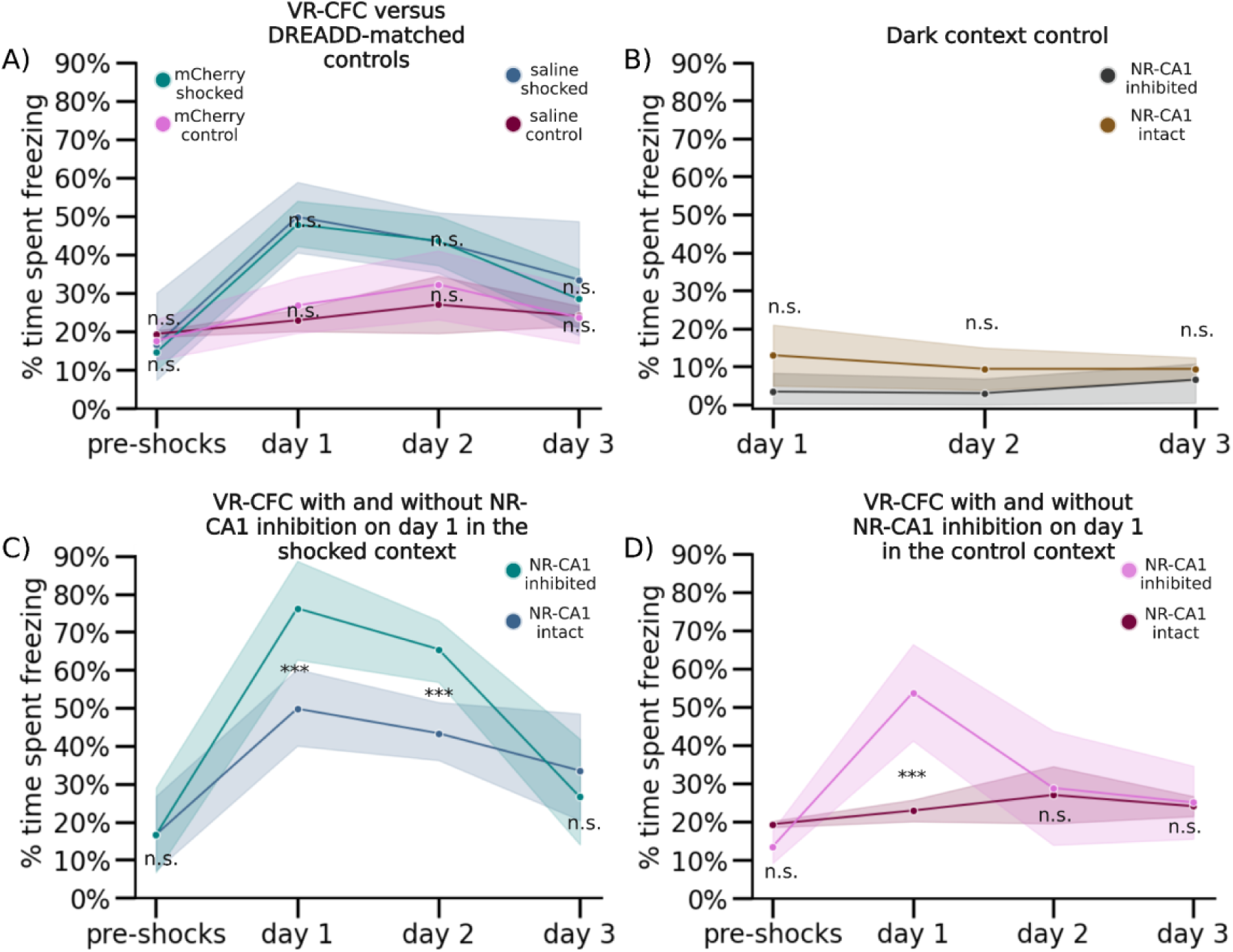
DREADD inhibition controls, context controls, and within context comparisons of NR-CA1 inhibition. (A) DREADD control mice were either injected with the hM4Di-lacking AAVrg-hSyn-DIO-mCherry (mCherry: see Method for details) and IP injected with 0.1mg/kg DCZ on day 1, 30 minutes before experimental start and quantity-matched saline on all other days, or injected with the h4MDi-intact AAVrg-hSyn-DIO-h4MDi-mCherry (Saline) and IP injected with saline on all days. No difference between the groups freezing activity was observed (Wilcoxon Rank Sum, p>0.05). (B) On retrieval days 1-3, we additionally recorded in a ‘dark’ context, a dark VR with no visual cues in NR-CA1 inactivated mice and DREADD control mice for the same length of time as context exposures. Mice froze at consistently low levels in the dark across days, with no difference in freezing levels between groups (Wilcoxon Rank Sum). (C) Direct comparison of freezing behavior in NR-CA1 inhibited mice (same data as Fig. 1E) versus uninhibited mice (DREADD control mice, same data as panel A) in the shocked context (Wilcoxon Rank Sum, P=day 1: 1.45e-5, day 2: 8.52e-3). (D) Same as C but in the control context (Wilcoxon Rank Sum, P=day 1: 9.58e-11).

**Extended Data Figure 4:**
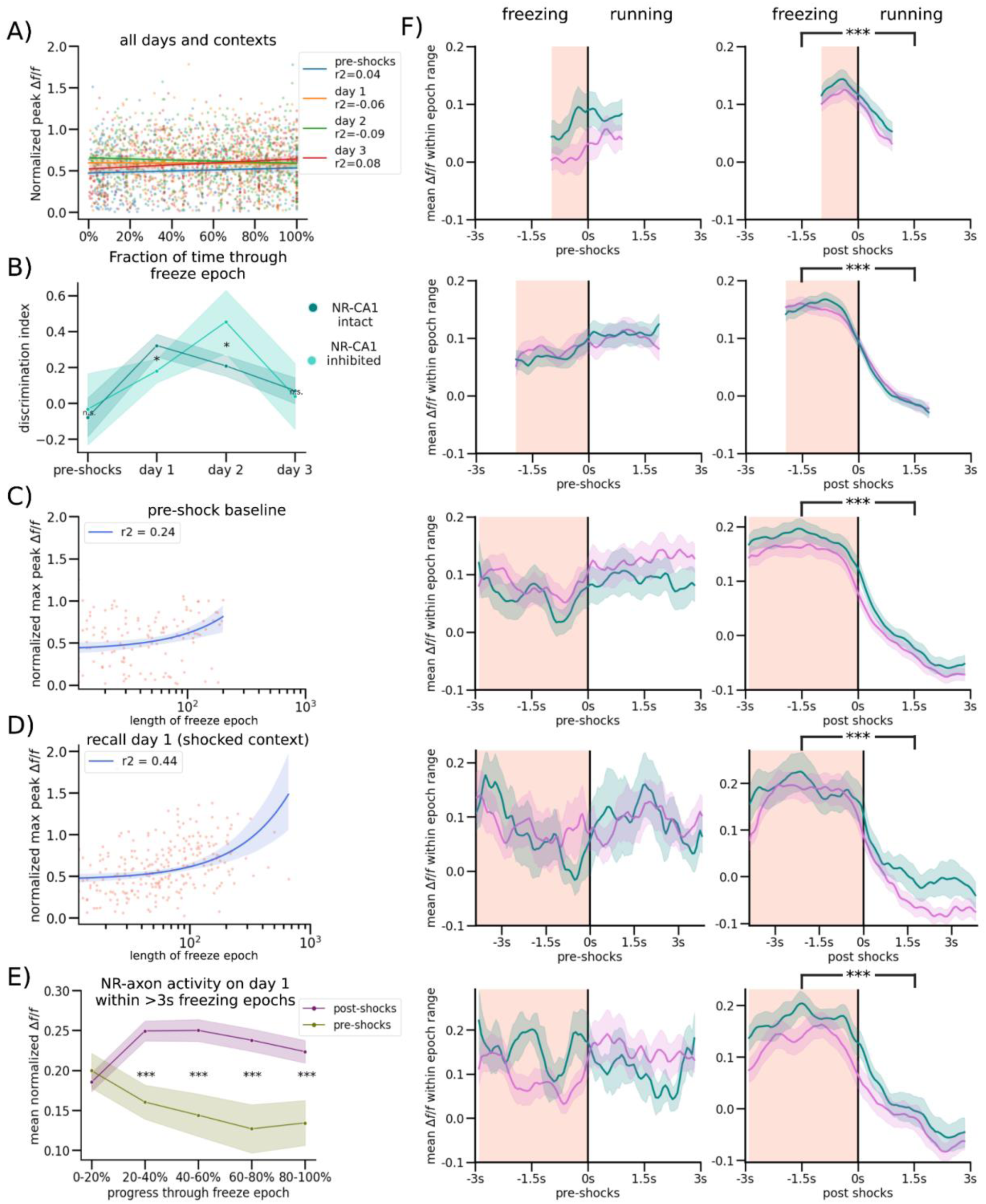
Context discrimination with and without NR-CA1 pathway inhibition and comparisons of NR-CA1 axonal activity before and after CFC. (A) To determine if there was a temporal aspect to the peak activity of each pause epoch, we plotted the max activity of each axon per freezing epoch on each day in both contexts (Dots, color-coded by experimental day), then plotted a robust linear regression regression to each day. This analysis indicated that the highest point of axonal activity within a freeze could occur anywhere temporally within a freezing epoch. (B) Discrimination index was calculated per day for both the NR-CA1 intact shocked mice and NR-CA1 DREADD inhibited shocked mice as (% time spent freezing in shocked context - % time spent freezing in control context)/total % time spent freezing in both contexts. Mice with an inhibited NR-CA1 pathway discriminated less between the two contexts under the effect of inhibition (Wilcoxon Rank Sum, P=0.0046), and discriminated better once the NR-CA1 pathway was uninhibited the following day (Wilcoxon Rank Sum, P=0.0033), than mice without inhibition, indicating an important role for an intact NR-CA1 pathway in discrimination following VR-CFC. (C) To test the impact of freeze length on maximum NR-CA1 axonal amplitude, we plotted the maximum peak within each freeze against the length of the freeze epoch in the NR-CA1 intact condition in the ‘shocked’ context before shocks (behavior plotted in Fig. 1D) with a robust quadratic regression (see Method Details for details) showing a slight tendency for increased maximum amplitude in longer freezing epochs: potentially caused by increased opportunities for freezing-related activity to take place in longer continuous freezing epochs.(D) Analysis is the same as in Extended Data Fig. 4C, but on retrieval day 1 in the shocked context. While the *r^2^* of the robust regression increased in the shocked context post-shocks compared to pre-shocks, since average freeze lengths also increased (as mice freeze for longer epochs post-shocks), it is difficult to assert that fearful freezing is inducing changes in normalized max axonal peak across all freezing epochs from these analyses. (E) Same analysis conducted as Fig. 2E, but on only a subset of longer pauses (3s+ in length) showing a similar average shape of activity throughout a freeze to Fig. 2E, with slightly higher activity towards the end of a freeze epoch, possibly due to the consistent NR-axonal activity ~0.5s before the end of a freezing epoch comprising a smaller fraction of the average time within the 80-100% of freeze bin. (F) Analyses conducted identically to Fig. 2D, except the freezing and post-freeze running epoch window was restricted to different window lengths of 1-2s, 2-3s, 4-5s, 5-6s, and 6-7s from top to bottom. All freezing epoch periods remained significantly elevated compared to running epoch periods post-shocks (Wilcoxon Rank Sum), with the same general underlying shape evident irrespective of the window chosen.

**Extended Data Figure 5:**
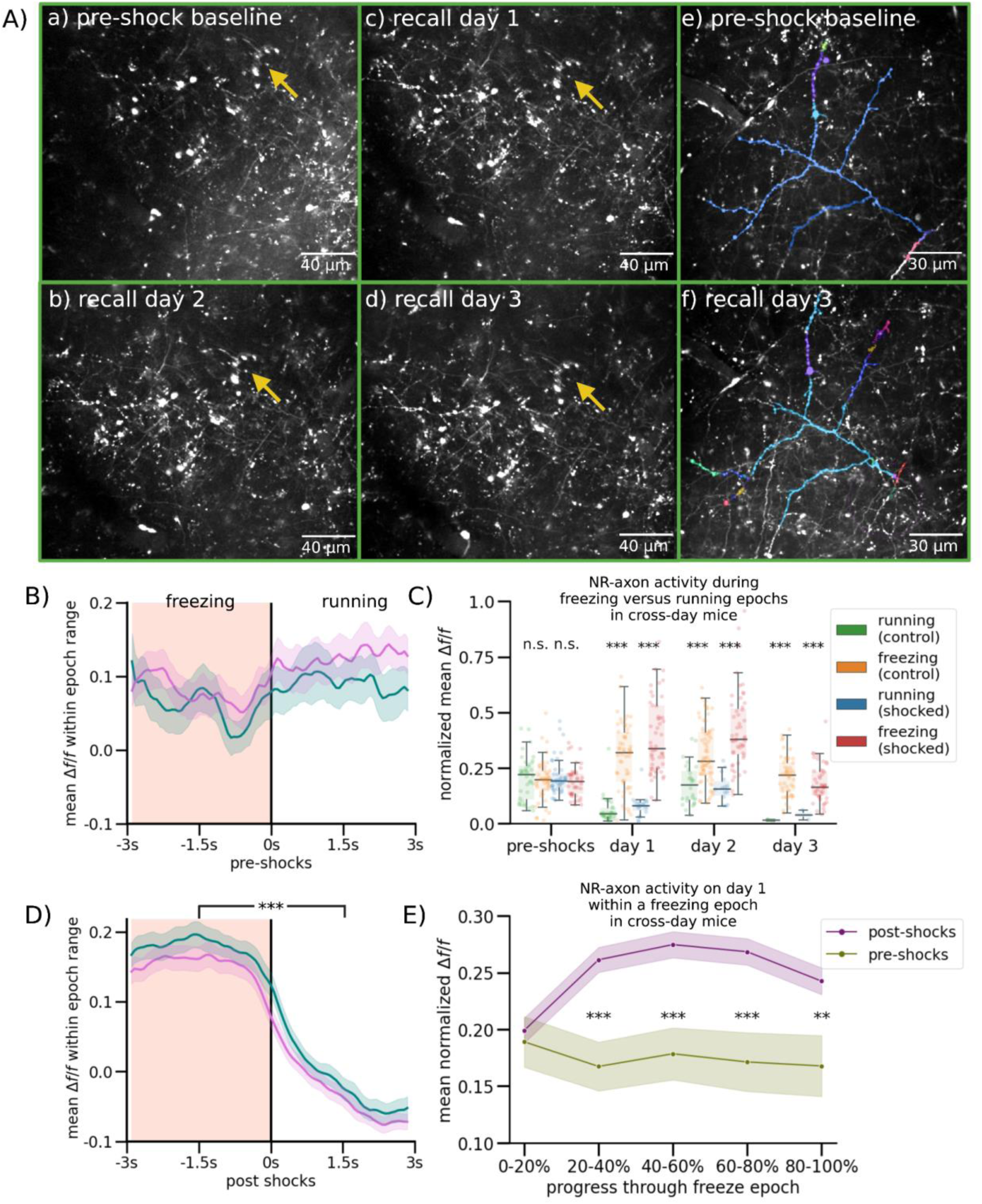
Multi-day tracking of the same NR-CA1 axons. (A) Two example fields of view are shown. All FOVs are directly outputted from the Suite2p mean image with no color correction modifications or crops applied. Left and middle column show a single FOV across three retrieval days, with yellow arrows pointing to the same structure over all four imaging days. Right column shows axonal structure from Fig. 2A (far right), with Suite2p ROIs that comprised the final combined trace highlighted on both the pre-shock baseline day and retrieval day 3, demonstrating our capacity to track the same NR-CA1 axonal structure over multiple days (N=4/10 imaged mice). (B) Analyses same as Fig. 2D (left), but on the subset of multi-day imaged axons. (C) Analyses same as Fig. 2C, but on the subset of multi-day imaged axons. (D) Analyses same as Fig. 2C (right), but on the subset of multi-day imaged axons. (E) Analyses same as Fig. 2E, but on the subset of multi-day imaged axons.

**Extended Data Figure 6:**
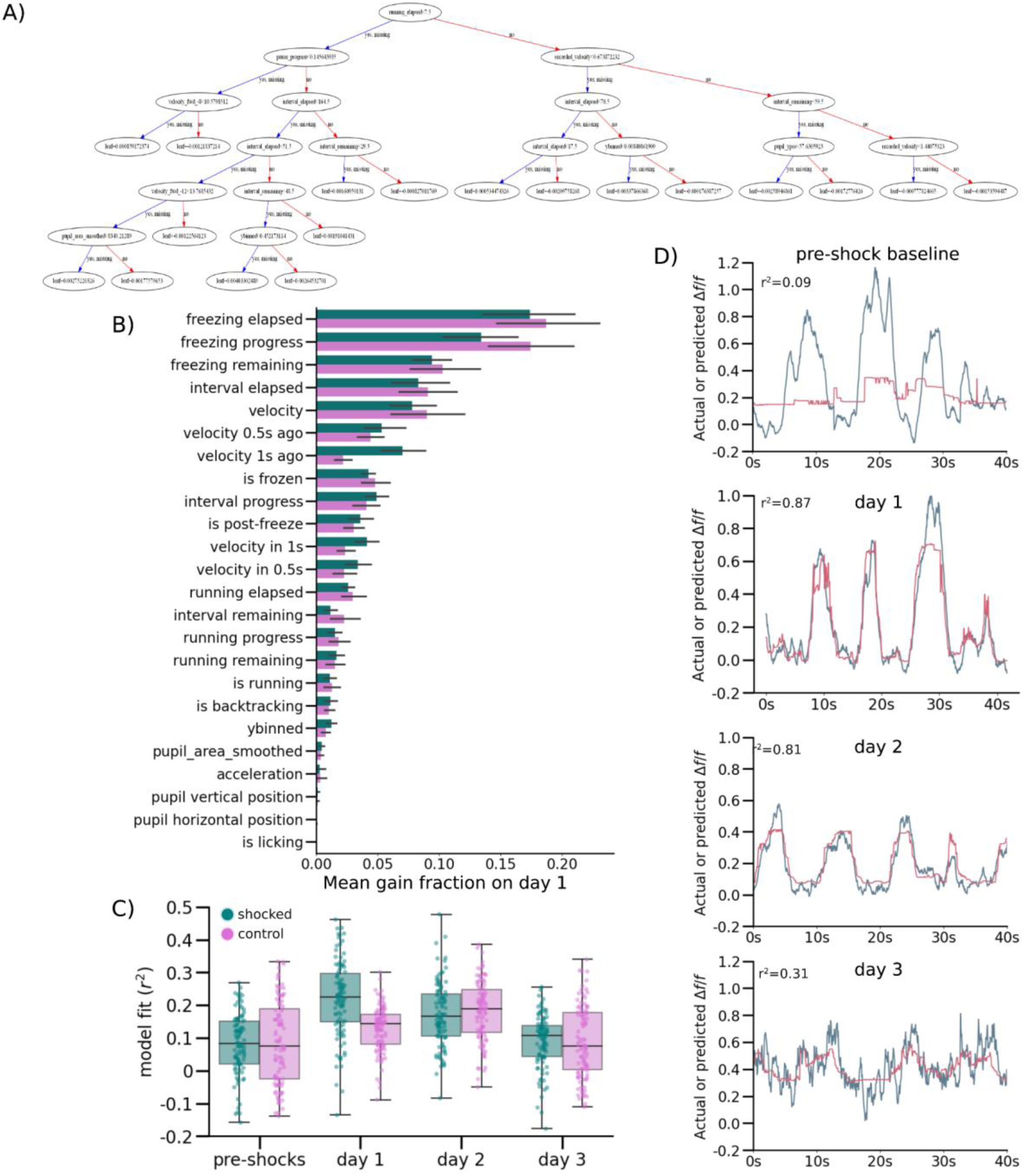
Computational Model Details and Additional Examples. (A) Real example of a model tree on retrieval day 1 in the feared context from the same example mouse shown in Fig. 3D. (B) Full ungrouped version of Fig. 3H. (C) Same analyses as Fig 3G, except from a model that ran without differentiating and building models per individual mouse, but instead across all mice and differentiating only on day and context, demonstrating that high inter-mouse variability in axonal activity and amplitude led to decreased median *r^2^* goodness-of-fit model performance compared to inter-mouse modeling shown in Fig. 3G. (D) Analyses same as Fig. 3C-F in the same mouse shown in Fig. 3C-F, but in the control instead of the shocked context.

## Notes

### Competing Interest Statement

The authors have declared no competing interest.

## References

1. Lissek, S., Kaczkurkin, A.N., Rabin, S., Geraci, M., Pine, D.S., and Grillon, C. (2014). Generalized anxiety disorder is associated with overgeneralization of classically conditioned fear. Biol. Psychiatry 75, 909–915.

2. Garfinkel, S.N., Abelson, J.L., King, A.P., Sripada, R.K., Wang, X., Gaines, L.M., and Liberzon, I. (2014). Impaired contextual modulation of memories in PTSD: an fMRI and psychophysiological study of extinction retention and fear renewal. J. Neurosci. 34, 13435–13443.

3. Kim, J.J., and Fanselow, M.S. (1992). Modality-specific retrograde amnesia of fear. Science 256, 675–677.

4. LeDoux, J.E. (2000). Emotion circuits in the brain. Annu. Rev. Neurosci. 23, 155–184.

5. Maren, S. (2001). Neurobiology of Pavlovian fear conditioning. Annu. Rev. Neurosci. 24, 897–931.

6. Maren, S., Phan, K.L., and Liberzon, I. (2013). The contextual brain: implications for fear conditioning, extinction and psychopathology. Nat. Rev. Neurosci. 14, 417–428.

7. Blanchard, R.J., and Blanchard, D.C. (1969). Crouching as an index of fear. J. Comp. Physiol. Psychol. 67, 370–375.

8. Fanselow, M.S., and Tighe, T.J. (1988). Contextual conditioning with massed versus distributed unconditional stimuli in the absence of explicit conditional stimuli. J. Exp. Psychol. Anim. Behav. Process. 14, 187–199.

9. Suzuki, A., Josselyn, S.A., Frankland, P.W., Masushige, S., Silva, A.J., and Kida, S. (2004). Memory reconsolidation and extinction have distinct temporal and biochemical signatures. J. Neurosci. 24, 4787–4795.

10. An, B., Kim, J., Park, K., Lee, S., Song, S., and Choi, S. (2017). Amount of fear extinction changes its underlying mechanisms. Elife 6. 10.7554/eLife.25224.

11. Kida, S. (2020). Function and mechanisms of memory destabilization and reconsolidation after retrieval. Proc. Jpn. Acad. Ser. B Phys. Biol. Sci. 96, 95–106.

12. Lee, C., Lee, B.H., Jung, H., Lee, C., Sung, Y., Kim, H., Kim, J., Shim, J.Y., Kim, J.-I., Choi, D.I., et al. (2023). Hippocampal engram networks for fear memory recruit new synapses and modify pre-existing synapses in vivo. Curr. Biol. 33, 507–516.e3.

13. Venkataraman, A., and Dias, B.G. (2022). Expanding the canon: An inclusive neurobiology of thalamic and subthalamic fear circuits. Neuropharmacology 226, 109380.

14. Frankland, P.W., Cestari, V., Filipkowski, R.K., McDonald, R.J., and Silva, A.J. (1998). The dorsal hippocampus is essential for context discrimination but not for contextual conditioning. Behav. Neurosci. 112, 863–874.

15. Guzowski, J.F., McNaughton, B.L., Barnes, C.A., and Worley, P.F. (1999). Environment-specific expression of the immediate-early gene Arc in hippocampal neuronal ensembles. Nat. Neurosci. 2, 1120–1124.

16. Wood, E.R., Dudchenko, P.A., Robitsek, R.J., and Eichenbaum, H. (2000). Hippocampal neurons encode information about different types of memory episodes occurring in the same location. Neuron 27, 623–633.

17. Hainmueller, T., and Bartos, M. (2018). Parallel emergence of stable and dynamic memory engrams in the hippocampus. Nature 558, 292–296.

18. Miry, O., Li, J., and Chen, L. (2020). The Quest for the Hippocampal Memory Engram: From Theories to Experimental Evidence. Front. Behav. Neurosci. 14, 632019.

19. Goode, T.D., Tanaka, K.Z., Sahay, A., and McHugh, T.J. (2020). An Integrated Index: Engrams, Place Cells, and Hippocampal Memory. Neuron 107, 805–820.

20. Matsuo, N. (2015). Irreplaceability of Neuronal Ensembles after Memory Allocation. Cell Rep. 11, 351–357.

21. Liu, X., Ramirez, S., Pang, P.T., Puryear, C.B., Govindarajan, A., Deisseroth, K., and Tonegawa, S. (2012). Optogenetic stimulation of a hippocampal engram activates fear memory recall. Nature 484, 381–385.

22. Ghandour, K., Ohkawa, N., Fung, C.C.A., Asai, H., Saitoh, Y., Takekawa, T., Okubo-Suzuki, R., Soya, S., Nishizono, H., Matsuo, M., et al. (2019). Orchestrated ensemble activities constitute a hippocampal memory engram. Nat. Commun. 10, 2637.

23. Vertes, R.P. (2002). Analysis of projections from the medial prefrontal cortex to the thalamus in the rat, with emphasis on nucleus reuniens. J. Comp. Neurol. 442, 163–187.

24. McKenna, J.T., and Vertes, R.P. (2004). Afferent projections to nucleus reuniens of the thalamus. J. Comp. Neurol. 480, 115–142.

25. Vertes, R.P. (2006). Interactions among the medial prefrontal cortex, hippocampus and midline thalamus in emotional and cognitive processing in the rat. Neuroscience 142, 1–20.

26. Vertes, R.P., Linley, S.B., and Hoover, W.B. (2015). Limbic circuitry of the midline thalamus. Neurosci. Biobehav. Rev. 54, 89–107.

27. Wolff, M., Alcaraz, F., Marchand, A.R., and Coutureau, E. (2015). Functional heterogeneity of the limbic thalamus: From hippocampal to cortical functions. Neurosci. Biobehav. Rev. 54, 120–130.

28. Dolleman-van der Weel, M.J., Griffin, A.L., Ito, H.T., Shapiro, M.L., Witter, M.P., Vertes, R.P., and Allen, T.A. (2019). The nucleus reuniens of the thalamus sits at the nexus of a hippocampus and medial prefrontal cortex circuit enabling memory and behavior. Learn. Mem. 26, 191–205.

29. Xu, W., and Südhof, T.C. (2013). A neural circuit for memory specificity and generalization. Science 339, 1290–1295.

30. Vetere, G., Kenney, J.W., Tran, L.M., Xia, F., Steadman, P.E., Parkinson, J., Josselyn, S.A., and Frankland, P.W. (2017). Chemogenetic Interrogation of a Brain-wide Fear Memory Network in Mice. Neuron 94, 363–374.e4.

31. Troyner, F., Bicca, M.A., and Bertoglio, L.J. (2018). Nucleus reuniens of the thalamus controls fear memory intensity, specificity and long-term maintenance during consolidation. Hippocampus 28, 602–616.

32. Ramanathan, K.R., Jin, J., Giustino, T.F., Payne, M.R., and Maren, S. (2018). Prefrontal projections to the thalamic nucleus reuniens mediate fear extinction. Nat. Commun. 9, 4527.

33. Ramanathan, K.R., Ressler, R.L., Jin, J., and Maren, S. (2018). Nucleus Reuniens Is Required for Encoding and Retrieving Precise, Hippocampal-Dependent Contextual Fear Memories in Rats. J. Neurosci. 38, 9925–9933.

34. Ramanathan, K.R., and Maren, S. (2019). Nucleus reuniens mediates the extinction of contextual fear conditioning. Behav. Brain Res. 374, 112114.

35. Totty, M.S., Ramanathan, K.R., Jin, J., Peters, S.E., and Maren, S. (2022). Thalamic nucleus reuniens coordinates prefrontal-hippocampal synchrony to suppress extinguished fear. bioRxiv, 2022.11.11.516165. 10.1101/2022.11.11.516165.

36. Moscarello, J.M. (2020). Prefrontal cortex projections to the nucleus reuniens suppress freezing following two-way signaled avoidance training. Learn. Mem. 27, 119–123.

37. Lovett-Barron, M., Kaifosh, P., Kheirbek, M.A., Danielson, N., Zaremba, J.D., Reardon, T.R., Turi, G.F., Hen, R., Zemelman, B.V., and Losonczy, A. (2014). Dendritic inhibition in the hippocampus supports fear learning. Science 343, 857–863.

38. Rajasethupathy, P., Sankaran, S., Marshel, J.H., Kim, C.K., Ferenczi, E., Lee, S.Y., Berndt, A., Ramakrishnan, C., Jaffe, A., Lo, M., et al. (2015). Projections from neocortex mediate top-down control of memory retrieval. Nature 526, 653–659.

39. Dong, C., Madar, A.D., and Sheffield, M.E.J. (2021). Distinct place cell dynamics in CA1 and CA3 encode experience in new environments. Nat. Commun. 12, 1–13.

40. Krishnan, S., Heer, C., Cherian, C., and Sheffield, M.E.J. (2022). Reward expectation extinction restructures and degrades CA1 spatial maps through loss of a dopaminergic reward proximity signal. Nat. Commun. 13, 6662.

41. Zhu, H., Pleil, K.E., Urban, D.J., Moy, S.S., Kash, T.L., and Roth, B.L. (2014). Chemogenetic inactivation of ventral hippocampal glutamatergic neurons disrupts consolidation of contextual fear memory. Neuropsychopharmacology 39, 1880–1892.

42. Roth, B.L. (2016). DREADDs for Neuroscientists. Neuron 89, 683–694.

43. Nagai, Y., Miyakawa, N., Takuwa, H., Hori, Y., Oyama, K., Ji, B., Takahashi, M., Huang, X.-P., Slocum, S.T., DiBerto, J.F., et al. (2020). Deschloroclozapine, a potent and selective chemogenetic actuator enables rapid neuronal and behavioral modulations in mice and monkeys. Nat. Neurosci. 23, 1157–1167.

44. Wouterlood, F.G., Saldana, E., and Witter, M.P. (1990). Projection from the nucleus reuniens thalami to the hippocampal region: light and electron microscopic tracing study in the rat with the anterograde tracer Phaseolus vulgaris-leucoagglutinin. J. Comp. Neurol. 296, 179–203.

45. Herkenham, M. (1978). The connections of the nucleus reuniens thalami: evidence for a direct thalamo-hippocampal pathway in the rat. J. Comp. Neurol. 177, 589–610.

46. Vertes, R.P., Hoover, W.B., Do Valle, A.C., Sherman, A., and Rodriguez, J.J. (2006). Efferent projections of reuniens and rhomboid nuclei of the thalamus in the rat. J. Comp. Neurol. 499, 768– 796.

47. Bertram, E.H., and Zhang, D.X. (1999). Thalamic excitation of hippocampal CA1 neurons: a comparison with the effects of CA3 stimulation. Neuroscience 92, 15–26.

48. Dolleman-Van der Weel, M.J., Lopes da Silva, F.H., and Witter, M.P. (1997). Nucleus reuniens thalami modulates activity in hippocampal field CA1 through excitatory and inhibitory mechanisms. J. Neurosci. 17, 5640–5650.

49. Dolleman-van der Weel, M.J., Lopes da Silva, F.H., and Witter, M.P. (2017). Interaction of nucleus reuniens and entorhinal cortex projections in hippocampal field CA1 of the rat. Brain Struct. Funct. 222, 2421–2438.

50. Goswamee, P., Leggett, E., and McQuiston, A.R. (2021). Nucleus Reuniens Afferents in Hippocampus Modulate CA1 Network Function via Monosynaptic Excitation and Polysynaptic Inhibition. Front. Cell. Neurosci. 15, 660897.

51. Chen, T., and Guestrin, C. (2016). XGBoost: A Scalable Tree Boosting System. arXiv [cs.LG].

52. Ji, J., and Maren, S. (2008). Differential roles for hippocampal areas CA1 and CA3 in the contextual encoding and retrieval of extinguished fear. Learn. Mem. 15, 244–251.

53. Lacagnina, A.F., Brockway, E.T., Crovetti, C.R., Shue, F., McCarty, M.J., Sattler, K.P., Lim, S.C., Santos, S.L., Denny, C.A., and Drew, M.R. (2019). Distinct hippocampal engrams control extinction and relapse of fear memory. Nat. Neurosci. 22, 753–761.

54. Silva, B.A., Astori, S., Burns, A.M., Heiser, H., van den Heuvel, L., Santoni, G., Martinez-Reza, M.F., Sandi, C., and Gräff, J. (2021). A thalamo-amygdalar circuit underlying the extinction of remote fear memories. Nat. Neurosci. 24, 964–974.

55. Goshen, I., Brodsky, M., Prakash, R., Wallace, J., Gradinaru, V., Ramakrishnan, C., and Deisseroth, K. (2011). Dynamics of retrieval strategies for remote memories. Cell 147, 678–689.

56. Corcoran, K.A., and Quirk, G.J. (2007). Activity in prelimbic cortex is necessary for the expression of learned, but not innate, fears. J. Neurosci. 27, 840–844.

57. Giustino, T.F., and Maren, S. (2015). The Role of the Medial Prefrontal Cortex in the Conditioning and Extinction of Fear. Front. Behav. Neurosci. 9, 298.

58. Bayer, H., and Bertoglio, L.J. (2020). Infralimbic cortex controls fear memory generalization and susceptibility to extinction during consolidation. Sci. Rep. 10, 1–13.

59. Thompson, B.M., Baratta, M.V., Biedenkapp, J.C., Rudy, J.W., Watkins, L.R., and Maier, S.F. (2010). Activation of the infralimbic cortex in a fear context enhances extinction learning. Learn. Mem. 17, 591–599.

60. Hoover, W.B., and Vertes, R.P. (2012). Collateral projections from nucleus reuniens of thalamus to hippocampus and medial prefrontal cortex in the rat: a single and double retrograde fluorescent labeling study. Brain Struct. Funct. 217, 191–209.

61. Steward, O., and Scoville, S.A. (1976). Cells of origin of entorhinal cortical afferents to the hippocampus and fascia dentata of the rat. The Journal of Comparative Neurology 169, 347–370. 10.1002/cne.901690306.

62. Kajiwara, R., Wouterlood, F.G., Sah, A., Boekel, A.J., Baks-te Bulte, L.T.G., and Witter, M.P. (2008). Convergence of entorhinal and CA3 inputs onto pyramidal neurons and interneurons in hippocampal area CA1--an anatomical study in the rat. Hippocampus 18, 266–280.

63. Chittajallu, R., Wester, J.C., Craig, M.T., Barksdale, E., Yuan, X.Q., Akgül, G., Fang, C., Collins, D., Hunt, S., Pelkey, K.A., et al. (2017). Afferent specific role of NMDA receptors for the circuit integration of hippocampal neurogliaform cells. Nat. Commun. 8, 152.

64. Sun, Y., Nguyen, A.Q., Nguyen, J.P., Le, L., Saur, D., Choi, J., Callaway, E.M., and Xu, X. (2014). Cell-type-specific circuit connectivity of hippocampal CA1 revealed through Cre-dependent rabies tracing. Cell Rep. 7, 269–280.

65. Andrianova, L., Brady, E.S., Margetts-Smith, G., Kohli, S., McBain, C.J., and Craig, M.T. (2021). Hippocampal CA1 pyramidal cells do not receive monosynaptic input from thalamic nucleus reuniens. bioRxiv, 2021.09.30.462517. 10.1101/2021.09.30.462517.

66. Morales, G.J., Ramcharan, E.J., Sundararaman, N., Morgera, S.D., and Vertes, R.P. (2007). Analysis of the actions of nucleus reuniens and the entorhinal cortex on EEG and evoked population behavior of the hippocampus. Conf. Proc. IEEE Eng. Med. Biol. Soc. 2007, 2480–2484.

67. Dolleman-Van Der Weel, M.J., and Witter, M.P. (1996). Projections from the nucleus reuniens thalami to the entorhinal cortex, hippocampal field CA1, and the subiculum in the rat arise from different populations of neurons. J. Comp. Neurol. 364, 637–650.

68. Sheffield, M.E.J., Adoff, M.D., and Dombeck, D.A. (2017). Increased Prevalence of Calcium Transients across the Dendritic Arbor during Place Field Formation. Neuron 96, 490–504.e5.

69. Sun, Q., Buss, E.W., Jiang, Y.-Q., Santoro, B., Brann, D.H., Nicholson, D.A., and Siegelbaum, S.A. (2021). Frequency-Dependent Synaptic Dynamics Differentially Tune CA1 and CA2 Pyramidal Neuron Responses to Cortical Input. J. Neurosci. 41, 8103–8110.

70. Milstein, A.D., Li, Y., Bittner, K.C., Grienberger, C., Soltesz, I., Magee, J.C., and Romani, S. (2021). Bidirectional synaptic plasticity rapidly modifies hippocampal representations. Elife 10. 10.7554/eLife.73046.

71. Priestley, J.B., Bowler, J.C., Rolotti, S.V., Fusi, S., and Losonczy, A. (2022). Signatures of rapid plasticity in hippocampal CA1 representations during novel experiences. Neuron 110, 1978–1992.e6.

72. Fan, L.Z., Kim, D.K., Jennings, J.H., Tian, H., Wang, P.Y., Ramakrishnan, C., Randles, S., Sun, Y., Thadhani, E., Kim, Y.S., et al. (2023). All-optical physiology resolves a synaptic basis for behavioral timescale plasticity. Cell 186, 543–559.e19.

73. Milstein, A.D., Li, Y., Bittner, K.C., Grienberger, C., Soltesz, I., Magee, J.C., and Romani, S. (2020). Bidirectional synaptic plasticity rapidly modifies hippocampal representations independent of correlated activity. 2020.02.04.934182. 10.1101/2020.02.04.934182.

74. Wang, M.E., Yuan, R.K., Keinath, A.T., Ramos Álvarez, M.M., and Muzzio, I.A. (2015). Extinction of Learned Fear Induces Hippocampal Place Cell Remapping. J. Neurosci. 35, 9122–9136.

75. Wu, C.-T., Haggerty, D., Kemere, C., and Ji, D. (2017). Hippocampal awake replay in fear memory retrieval. Nat. Neurosci. 20, 571–580.

76. Broussard, G.J., Liang, Y., Fridman, M., Unger, E.K., Meng, G., Xiao, X., Ji, N., Petreanu, L., and Tian, L. (2018). In vivo measurement of afferent activity with axon-specific calcium imaging. Nat. Neurosci. 21, 1272–1280.

77. Krashes, M.J., Koda, S., Ye, C., Rogan, S.C., Adams, A.C., Cusher, D.S., Maratos-Flier, E., Roth, B.L., and Lowell, B.B. (2011). Rapid, reversible activation of AgRP neurons drives feeding behavior in mice. J. Clin. Invest. 121, 1424–1428.

78. Schindelin, J., Arganda-Carreras, I., Frise, E., Kaynig, V., Longair, M., Pietzsch, T., Preibisch, S., Rueden, C., Saalfeld, S., Schmid, B., et al. (2012). Fiji: an open-source platform for biological-image analysis. Nat. Methods 9, 676–682.

79. Pachitariu, M., Stringer, C., Schröder, S., Dipoppa, M., Federico Rossi, L., Carandini, M., and Harris, K.D. (2016). Suite2p: beyond 10,000 neurons with standard two-photon microscopy. bioRxiv, 061507. 10.1101/061507.

80. Dombeck, D.A., Harvey, C.D., Tian, L., Looger, L.L., and Tank, D.W. (2010). Functional imaging of hippocampal place cells at cellular resolution during virtual navigation. Nat. Neurosci. 13, 1433–1440.

81. Aronov, D., Nevers, R., and Tank, D.W. (2017). Mapping of a non-spatial dimension by the hippocampal–entorhinal circuit. Nature 543, 719.

82. Kaufman, A.M., Geiller, T., and Losonczy, A. (2020). A Role for the Locus Coeruleus in Hippocampal CA1 Place Cell Reorganization during Spatial Reward Learning. Neuron 105, 1018–1026.e4.

